# Analysis of Transcriptograms in Epithelial-Mesenchymal Transition (EMT)

**DOI:** 10.64898/2026.02.16.706231

**Authors:** Odilon Júlio dos Santos, Rodrigo Juliani Siqueira Dalmolin, Rita M. C. de Almeida

## Abstract

Single-cell RNA sequencing (single-cell RNA-seq) has represented a revolution in gene expression analysis. However, high dropout rates and stochastic noise often reduce the amount of information captured in these experiments. The epithelial–mesenchymal transition (EMT), which is fundamental to tumor progression and organismal development, is particularly difficult to fully characterize due to the existence of intermediate states. In this work, we demonstrate that projecting transcriptomic data onto gene lists ordered using protein–protein interaction (PPI) information acts as a “biological low-pass filter”, attenuating technical noise and increasing the statistical power of the analyses.

We propose and validate an innovative pipeline that integrates the Transcriptogram method with Principal Component Analysis (PCA). By applying a moving average over functionally ordered genes, we drastically increase the signal-to-noise ratio, enabling the inference of cellular trajectories. The method was applied to a public dataset of TGF-*β*1–induced MCF10A cells, with rigorous batch-effect correction based on biological controls.

The results reveal that EMT is not merely a morphological change, but a coordinated, systemic reprogramming. This approach enabled the identification of critical modules that would remain hidden in conventional analyses: (i) a massive “Metabolic Switch” (Cluster 2), indicating a transition toward oxidative phosphorylation to sustain invasion; (ii) a strategic blockade of the cell cycle (Cluster 4); and (iii) a “Detoxification Shield” and chemoresistance program (Cluster 5), characterized by endogenous activation of metallothioneins.

We conclude that the combination of PPI network topology and dimensionality reduction offers superior resolution for dissecting cellular plasticity. The method not only validates classical markers, but also reveals the hidden functional architecture of the transition, showing that EMT is not a single, uniform process, but rather one in which cells can follow distinct trajectories, halting at different stages of differentiation.

## I. Introduction

**L**IFE arises from an extremely complex, nonlinear network of reactions involving numerous components, such as proteins, RNA, and metabolites within the cell. A cell’s current metabolic state influences its future metabolic configuration; when these states remain sufficiently similar over time, the cell preserves its identity. In dynamical terms, such a condition can be interpreted as a fixed point of the cell’s biochemical dynamics.

Given this remarkable complexity, accurately characterizing a cell state is a major challenge. One practical and effective approach has been the identification of molecular markers—a relatively small set of genes whose expression patterns distinguish specific cellular phenotypes. Markers associated with cancer, different stages of the cell cycle, or epithelial and mesenchymal phenotypes, for instance, are widely and successfully used to define and classify cell states.

However, molecular markers are often insufficient to reliably predict a cell’s future behavior. Cancer provides a paradigmatic example: even when a population of cancer cells expresses the same set of markers, substantial heterogeneity is still observed. This variability arises partly from stochastic external stimuli, but also from an incomplete characterization of the cell’s internal molecular state.

Because biochemical reaction networks are highly nonlinear, they may exhibit strong sensitivity to initial conditions. As a result, future biochemical states can diverge significantly, even when the initial states were classified as identical based on limited or coarse molecular profiling.

In this work, we introduce a method for analyzing single-cell RNA-seq data that leverages the richness of these datasets while applying statistical approaches to selectively discard information that is not relevant to our objectives. Noise is an inherent challenge in transcriptomic analyses. To mitigate its impact, we employ transcriptograms [1], [2], which are smoothed gene expression profiles that enhance the signal-to-noise ratio in gene expression measurements [3] (see the Methodology section below).

These transcriptograms are then used as input for Principal Component Analysis (PCA). Each principal component represents an expression profile in its own right, and the observed profiles can be reconstructed as linear combinations of these components. When most of the variance in the data is captured by a small number of principal components, the effective dimensionality of the problem is substantially reduced.

We applied this approach to the study of Epithelial–Mesenchymal Transition (EMT), using publicly available data generated by Deshmukh and collaborators [4]. EMT is a complex biological program in which epithelial cells downregulate adhesion complexes, lose apical–basal polarity, and acquire a mesenchymal phenotype characterized by increased migratory and invasive properties. Although EMT is essential for morphogenesis, its aberrant reactivation in neoplastic contexts is closely associated with metastasis and therapeutic resistance [5].

EMT was historically described as a binary process. However, single-cell RNA sequencing (scRNA-seq) studies have reshaped this perspective, revealing a dynamic continuum of intermediate states [4]. These findings highlight that the conventional set of marker genes is insufficient to fully characterize EMT, and that analyzing these transitional states remains computationally demanding. Moreover, standard dimensionality reduction methods typically treat genes as independent variables, overlooking the biological fact that genes function within highly interconnected regulatory and functional networks. The transcritogram method provides an automatic way to consider correlation between genes that belong to the same pathway or Gene Ontology term.

The main contributions of this paper are summarized as follows:

- We adapt the Transcriptogram methodology to single cell RNA-Seq, providing a **Biological Low-Pass Filter** approach that utilizes PPI network topology to smooth scRNA-seq data, reducing noise and enhancing signal-to-noise ratio.
- We demonstrate that the **Transcriptogram-based dimensionality reduction** achieves faster variance saturation (elbow point) compared to raw data, allowing for a more efficient reconstruction of cellular trajectories.
- We obtain a differential gene expression profile, associated to the first principal component provided by the PCA, whose presence in the samples serves as a proxy for EMT progression. As this profile is genome wide, it may provide a genome-wide marker profile for EMT. As we have only applied for one dataset we cannot claim it is universal. We intend to explore this further in the future.
- We uncover a **Metabolic Switch** (Oxidative Phosphorylation) and a **Detoxification Shield** (Metallothioneins) as intrinsic components of the EMT program, validating the method’s capability to discover non-canonical functional modules.
- We provide a reproducible computational pipeline that integrates topological smoothing, trajectory reconstruction, and functional “rescue” of gene clusters, applicable to other cellular plasticity models.

## II. Methodology

### A. Computational Pipeline Overview

The analytical framework developed in this study combines network-based noise attenuation with dimensionality reduction to analyze single-cell transcriptomic data. As outlined in Fig. 1, the workflow begins with raw data acquisition and stringent quality control, followed by application of the Transcriptogramer algorithm [1], [6]. The processed data are then subjected to PCA decomposition, enabling subsequent functional characterization of the resulting gene modules.

**Fig. 1.**
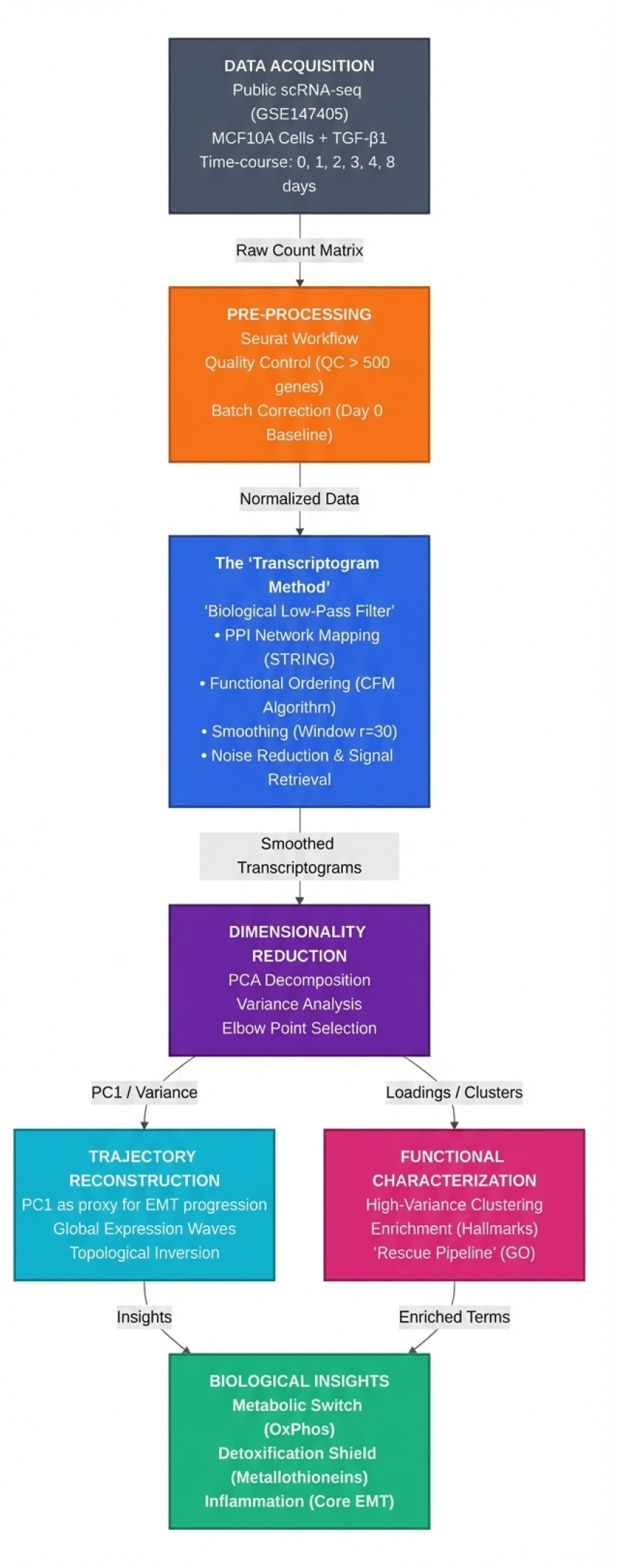
Integrative Computational Pipeline. The flowchart details the sequential steps: (1) Pre-processing of raw scRNA-seq data; (2) The core *Transcriptogramer* method acting as a biological low-pass filter; (3) PCA-based dimensionality reduction and trajectory reconstruction; (4) Functional clustering; and (5) The resulting biological insights regarding metabolism and inflammation.

### B. Data Acquisition and Batch Correction

Publicly available scRNA-seq data from MCF10A cells treated with TGF-*β*1 were obtained from [4]. The dataset comprises two batches of cells. The first one consists of samples treated up to 0, 4, and 8 days. The second batch consists of samples treated up to 0, 1, 2, and 3 days. The batches were processed in two different platforms (10x Genomics v2 and v3). Quality control was performed using the Seurat R package [7], filtering cells with fewer than 500 detected genes or mitochondrial gene content greater than 20%.

To mitigate technical variability between the two experimental batches (Batch 1 and Batch 2), we applied a linear scaling correction based on the biological controls (Day 0 samples), which serve as a common baseline. Since both batches contain untreated cells representing the same biological state, systematic deviations in their mean gene expression profiles were attributed to batch effects.

For each gene *g*, we calculated the mean expression 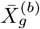 for each batch *b* using exclusively the Day 0 cells. We then defined a scaling factor *K*_*g*_ as the ratio between the mean expression in the reference batch (Batch 2) and the target batch (Batch 1):

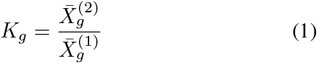

The normalized expression values *X*_*g,c*_ for all cells *c* in Batch 1 were then adjusted as:

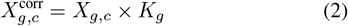

This linear transformation aligns the basal expression levels of the controls across datasets while preserving the relative biological variation induced by the TGF-*β*1 treatment in subsequent time points. The correction was implemented in R using vectorized operations.

### C. The Transcriptogram Method

The central component of our methodology is the Transcriptogramer package [1], [6]. In contrast to conventional approaches that consider genes as independent variables, this method projects gene expression data onto a gene list that has been ordered using information from Protein-Protein Interaction (PPI) networks (STRING-DB, score ≥ 800).

#### 1) Functional Ordering (CFM Algorithm)

The linear ordering of genes is not arbitrary. It is obtained by minimizing a cost function *C* that penalizes the distance between interacting proteins in the 1D list:

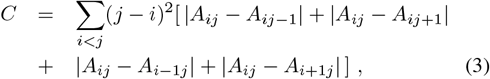

where *A*_*ij*_ is the adjacency matrix of the PPI network. This optimization is performed using a Monte Carlo algorithm (Simulated Annealing), ensuring that functionally related genes are placed in close proximity [1].

#### 2) Biological Low-Pass Filter

We applied a moving average window of radius *r* = 30 (encompassing 61 genes) to smooth the expression profiles over this ordered list. This process acts as a **biological low-pass filter**:

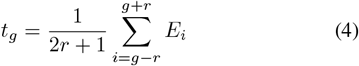

where *E*_*g*_ is the expression of the gene at position *g*, where *t*_*g*_ denotes the transcriptogram value corresponding to that gene. This smoothing procedure reduces stochastic noise (high-frequency fluctuations) inherent to scRNA-seq data while enhancing signals associated with coordinated functional modules (low-frequency variation) [3].

Notably, setting *r* = 0 yields transcriptograms in which raw expression values are assigned to each gene, without any smoothing. Even in this case, however, the result is not identical to the original expression matrix, since we restrict the analysis to genes included in the ordered list—namely, those whose protein products are recorded in the STRING database as having at least one protein–protein interaction with a score of 800 or higher. This preliminary filtering step reduces the number of genes from approximately 30,000 to about 14,000.

### D. PCA and Trajectory Reconstruction

Principal Component Analysis (PCA) was applied to the smoothed transcriptograms to achieve dimensionality reduction. Transcriptomes were then reconstructed using a subset of principal components (PCs), and the reconstruction error was evaluated by comparing the reconstructed transcriptograms with the original ones. The expression for reconstructing the transcriptome of a cell *c* using the first *k* components is given by:

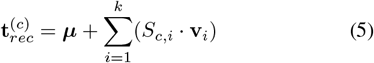

where ***µ*** and 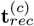 are, respectively, the mean expression vector and the reconstructed transcriptogram vector, *S*_*c,i*_ is the coefficiet of cell *c* in the durection defined by PC *i*, and **v**_*i*_ is the *i*th principal component. This step allows for the visualization of global “waves” of gene expression along the EMT progression.

### F. Functional Characterization and “Rescue” Pipeline

To interpret the biological patterns, we identified regions of high variance between samples along the transcriptograms. We then clustered them into functional modules using k-means. These clusters were initially characterized using the MSigDB Hallmark collection via the `clusterProfiler` package.

To overcome the limited coverage of general Hallmarks, we implemented a **“Rescue Pipeline”**. For clusters showing no significant Hallmark enrichment (e.g., Cluster 5), we automatically queried the Gene Ontology (GO) Biological Processes database. This hierarchical approach allowed us to identify specific mechanisms, such as “Response to Metal Ion” and “Detoxification”, which are critical for cell survival but often overlooked in EMT standard broad-spectrum analyses.

## III. Results

### A. Method Validation: Dimensionality Reduction Efficiency

Before dissecting the biological mechanisms, we evaluated the capacity of the Transcriptogramer method using *r* = 30 (R30 samples) to compress relevant biological information.

However, the smoothing step is decisive for signal consolidation. As shown in **Fig. 2**, the variance decomposition analysis of the R30 transcriptograms reveals a rapid accumulation of the explained variance. The detailed view of the first 25 components **(Left Panel)** clearly identifies the “elbow point” (saturation of the relevant topological signal), demonstrating that the core biological trajectory is captured within the first few dimensions. The global spectrum **(Right Panel)** confirms that higher-order components contribute marginally, mostly representing stochastic noise. This efficiency justifies the use of a low-dimensional space for the trajectory reconstruction in the subsequent analyses.

**Fig. 2.**
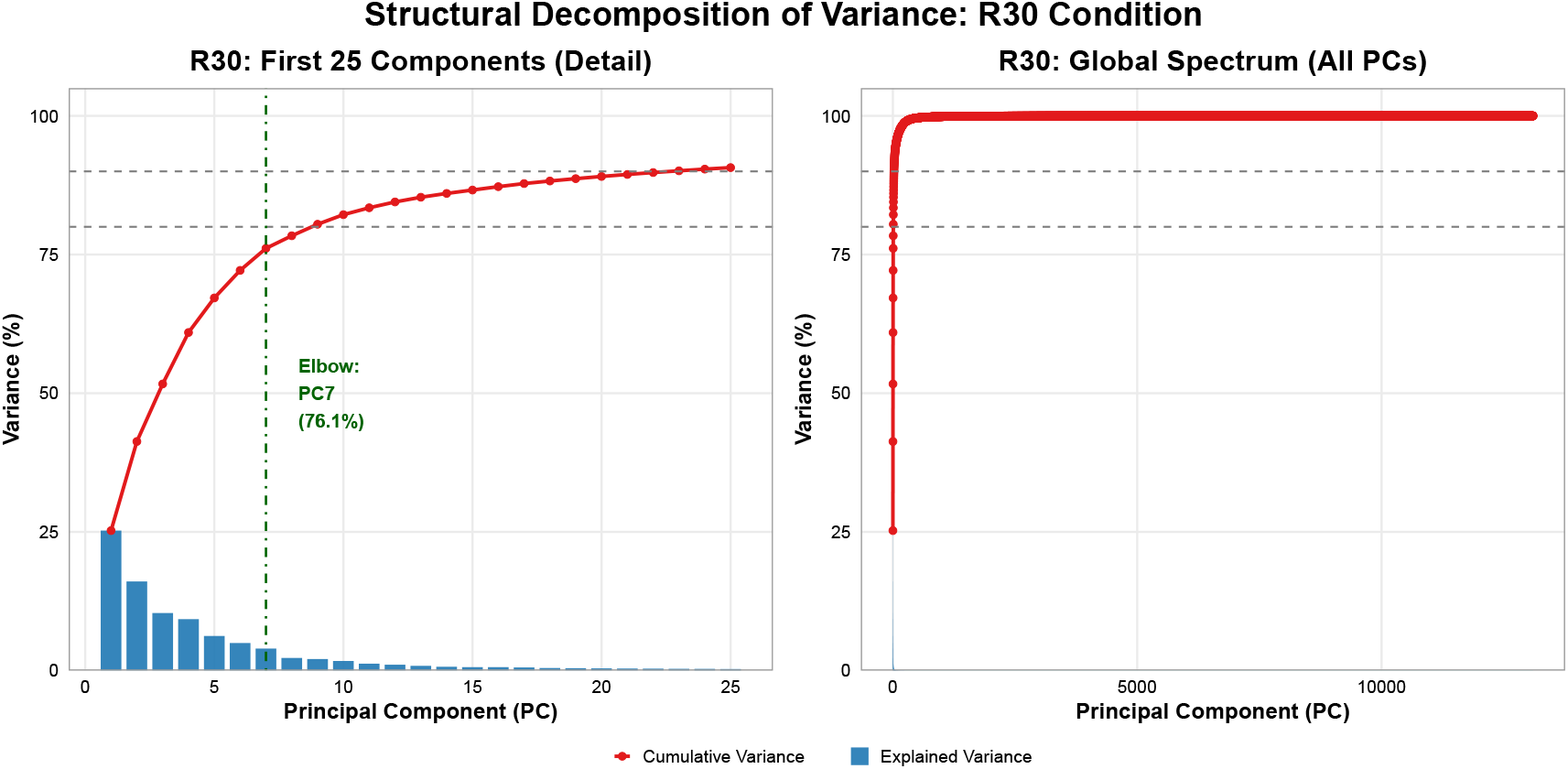
Structural Decomposition of Variance in R30 Condition. **(Left)** Detailed analysis of the first 25 Principal Components highlights the “elbow point,” indicating where the accumulation of biological signal saturates. **(Right)** The global spectrum (all components) shows the long tail of low-variance dimensions, confirming that topological smoothing effectively compresses the system’s dynamics into the leading components, filtering out noise.

### B. Trajectory Reconstruction

Fig. 3 presents the plots of the values of coefficients of several principal components as a function of the coefficients of the first principal component, PC1.

**Fig. 3.**
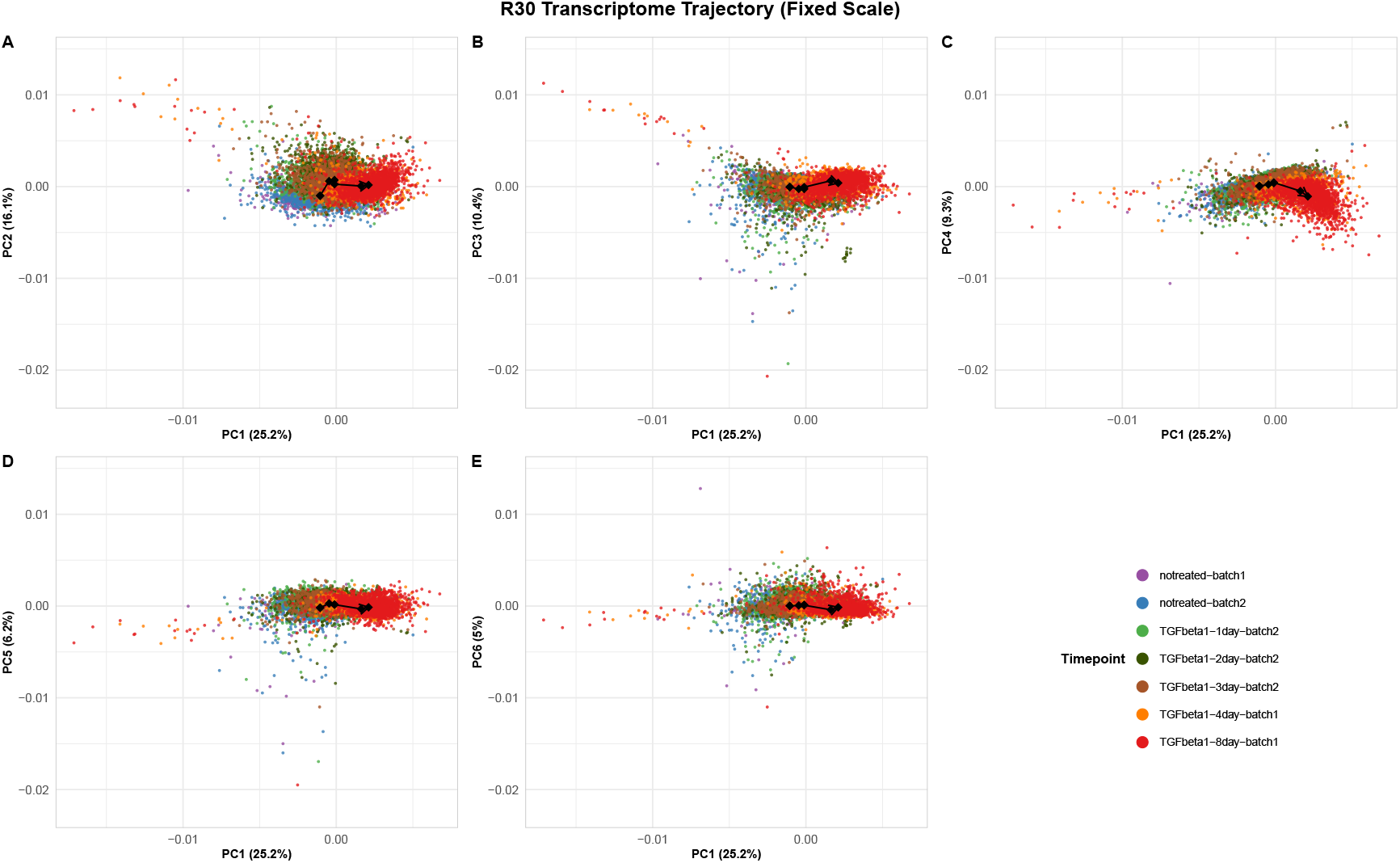
Structural Decomposition of Variance and Trajectory Analysis (R30). Plots of higher-order components coefficients (PC2 to PC6) against first Principal Component coefficient (PC1). The black arrows connect the centroids of each time point, illustrating the trajectory of the system’s center of mass from Day 0 to Day 8. **(A-C, Top Row)** PC2, PC3, and PC4 vs PC1 reveal a robust temporal structure, with clear separation between early (purple/blue), intermediate (green), and late (orange/red) stages. **(D-E, Bottom Row)** As dimensionality increases (PC5 and PC6), the temporal segregation decreases, confirming that the main biological signal driving the EMT is concentrated in the first few dimensions. The color legend (bottom right) applies to all panels.

### C. Global Dynamics: Waves of Expression

We first plot in Fig. 2 the values of coeficients relative to different components (PC2 up to PC6) as a function of the coeficient for the first principal component PC1 for all the samples, in different collors for each day. As shown in Fig. 1, upper, left panel, components 1 to 6 concentrate more than 75% of the total variance. There is a clear progression from smaller to larger values of PC1 coeficient for samples harvested at later days, showing that we can use this parameter as a proxy for EMT progression. The resulting analysis reveals that EMT does not occur as an abrupt switch, but as a pattern of “expression waves” as the cells progress in EMT (Fig. 4). This is useful for further analyses.

**Fig. 4.**
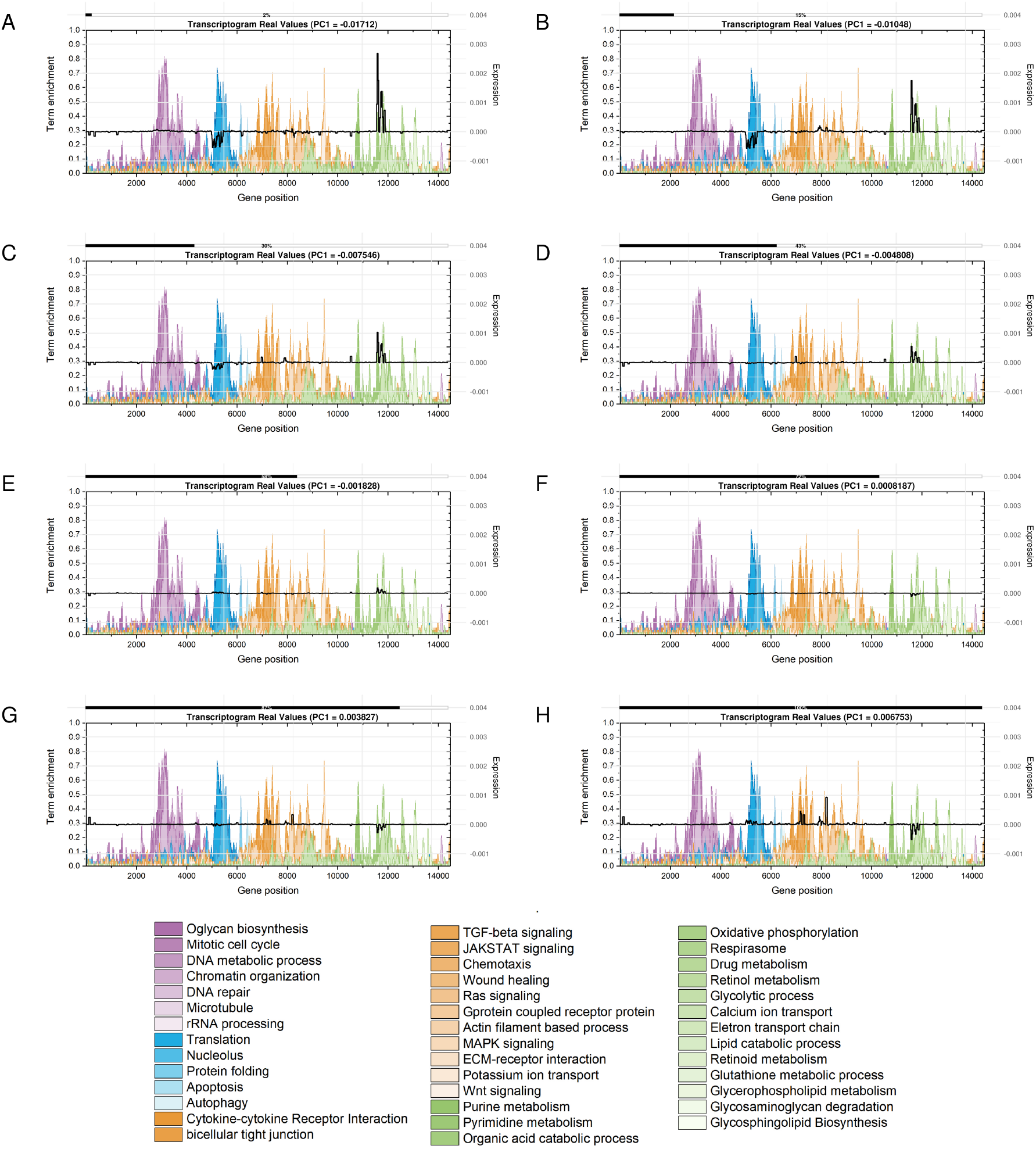
Evolutionary Panel of Global Transcriptome Reconstruction. The sequence of panels illustrates the topological dynamics of gene expression waves along the ordered gene list. Valley regions (low expression) in early phases progressively transform into peaks (high expression) in later phases. The color bar (bottom) indicates the relative expression levels. **(A)** 0% - Epithelial State; **(B)** 14% - Initiation; **(C)** 28% - Early Transition; **(D)** 42% - Plasticity Peak I; **E)** 57% - Plasticity Peak II; **(F)** 71% - Late Transition; **(G)** 85% - Mesenchymal Est.; **(H)** 100% - Mesenchymal Final.

To visualize the continuous nature of the transition, we reconstructed all transcriptomes using 6 principal components (from PC1 to PC6). Then, we subdivided the samples by their PC1 values into 50 intervals.. For the samples localized at the same PC1 coefficient interval, we calculated the average transcriptograms. This is shown in Fig **Fig. 4**, for different stages of EMT progression. The transcriptograms in this figure represents the expression changes in respect to the global transcriptogram average, or the center of mass of the points representing the samples in the expression space used in PCA. Hence, zero value in these transcriptograms represents zero change in respect to the average sample. As background, these figures show pathways or GO terms localization on the gene ordering. This is meant to help the biological interpretation of the observed gene expression alterations.

### D. Dynamic Orchestration of Functional Modules

The gene variance detection algorithm segmented the transcriptogram into co-regulated gene clusters. To understand the temporal hierarchy of these modules, we analyzed their relative contribution to the global variance along the EMT progression.

Fig. 5 illustrates this “changing of the guard”. We observe a sequential orchestration where modules responsible for epithelial maintenance decline, giving way to modules driving the mesenchymal phenotype. Crucially, the normalized importance plot (Fig. 5 TOP) and the stacked area plot (Fig. 5 BOTTOM) reveal that the transition is supported by a massive metabolic shift (Cluster 2) that persists and expands towards the end of the trajectory.

**Fig. 5.**
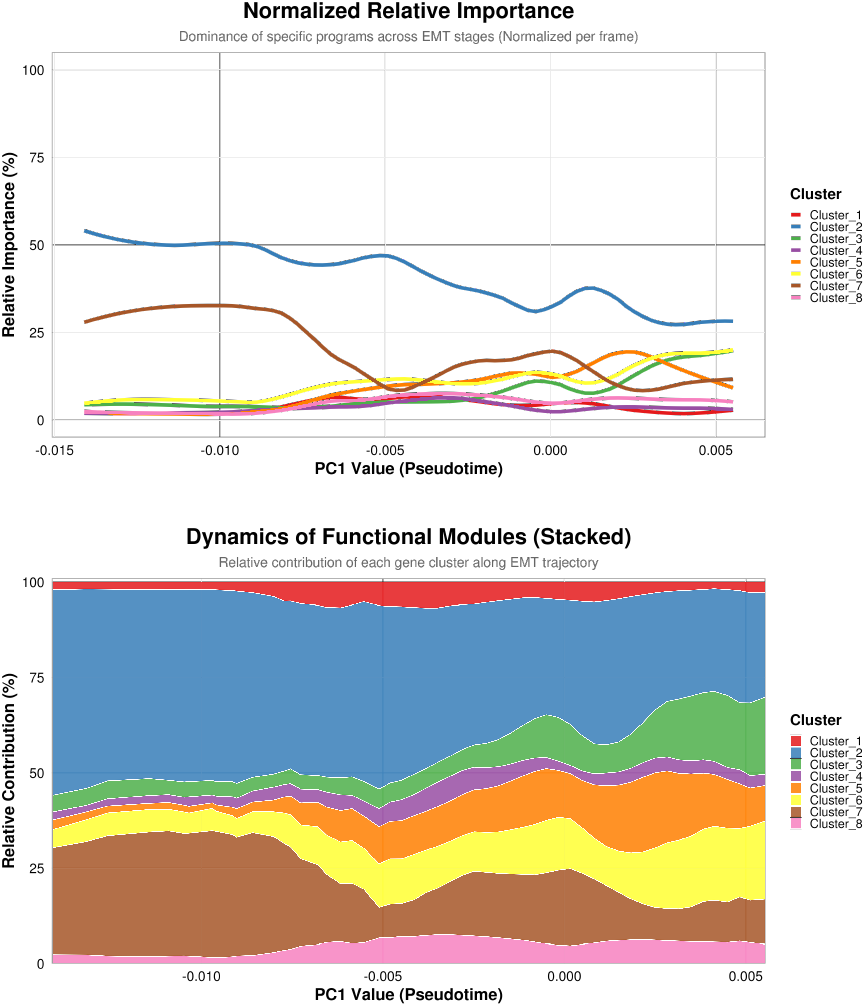
Dynamics of Functional Modules. **(Top)** Normalized Relative Importance plot showing the shifting dominance of specific gene clusters along the pseudotime. **(Bottom)** Stacked Contribution Area plot confirming that EMT is a continuous replacement of genetic programs, with no periods of transcriptional silence. Note the expansion of Cluster 2 (Metabolism) and Cluster 6 (Core EMT/Inflammation) in the later stages.

### E. Characterization of Key Modules

#### 1) The “Core” EMT and Inflammation (Cluster 6)

Cluster 6 was identified as the central driver, showing robust enrichment for the *Epithelial Mesenchymal Transition* signature (*p* < 10^*−*19^) and inflammatory pathways (*TNFA signaling via NFKB*), suggesting inflammation is intrinsic to the mesenchymal state (Fig. 6).

**Fig. 6.**
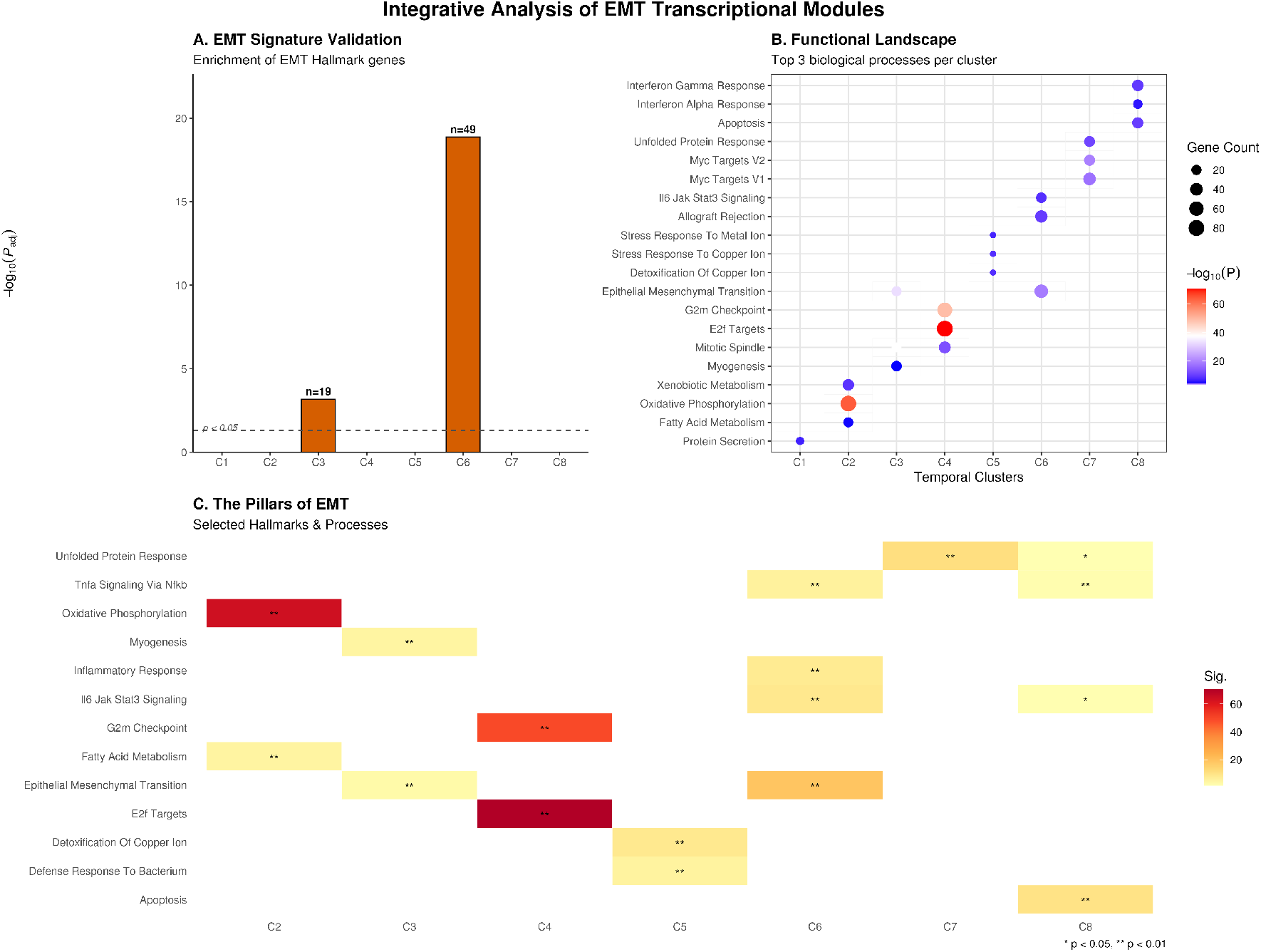
Functional Architecture of EMT. (A) Statistical validation of EMT signature in Clusters 3 and 6. (B) Global landscape revealing parallel processes. (C) Heatmap of the functional pillars: Plasticity, Bioenergetics, Inflammation, and Stress

#### 2) The Metabolic Switch (Cluster 2)

Cluster 2 represents a purely metabolic module, massively enriched for *Oxidative Phosphorylation* (*p* < 10^*−*64^). This confirms the bioenergetics reprogramming required to sustain the high energy demands of invasion.

#### 3) The Detoxification Shield (Cluster 5)

Cluster 5 was rescued by our Gene Ontology pipeline, revealing functions related to *Response to Metal Ion* and *Detoxification*. The upregulation of metallothioneins (*MT1X, MT2A*) indicates an endogenous chemoresistance mechanism activated during the transition.

### F. Comparative Analysis and Population Heterogeneity

We compared our findings with canonical markers described by Deshmukh et al. [4]. As shown in Fig. 7, while there is convergence on structural markers, our network-based approach uniquely highlights the Metabolic Switch and Defense Shield.

**Fig. 7.**
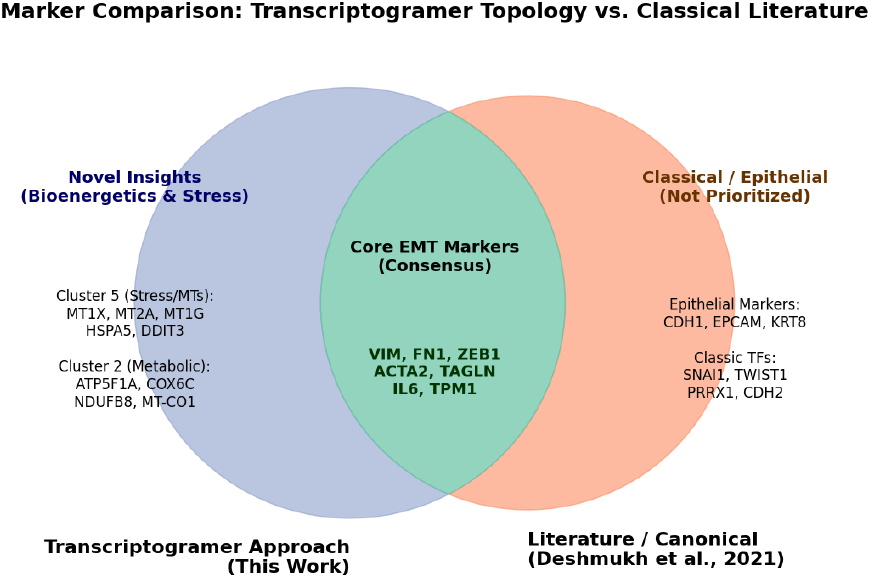
Comparative Analysis of Gene Scope: Transcriptogramer vs. Literature. Overlap analysis between the topological clusters identified in this study and the classical EMT markers (Deshmukh et al., 2021). (1) **Consensus:** Core EMT effectors were successfully retrieved, confirming method robustness. (2) **Novel Insights:** Our approach uniquely prioritized mitochondrial (Cluster 2) and stress-response genes (Cluster 5), suggesting that metabolic reprogramming and oxidative stress management are integral components of the EMT program. (3) **Literature-Exclusive:** Markers such as *CDH1* (epithelial) were excluded by the topological filter as they represent the downregulated state rather than the active mesenchymal transition.

Finally, analysis at the experimental endpoint (Day 8) revealed marked heterogeneity (Fig. 8). A significant fraction of cells did not reach the mesenchymal extreme, confirming the existence of stable hybrid phenotypes. This persistent hybrid population creates a reservoir of plasticity potentially linked to tumor dormancy or relapse.

**Fig. 8.**
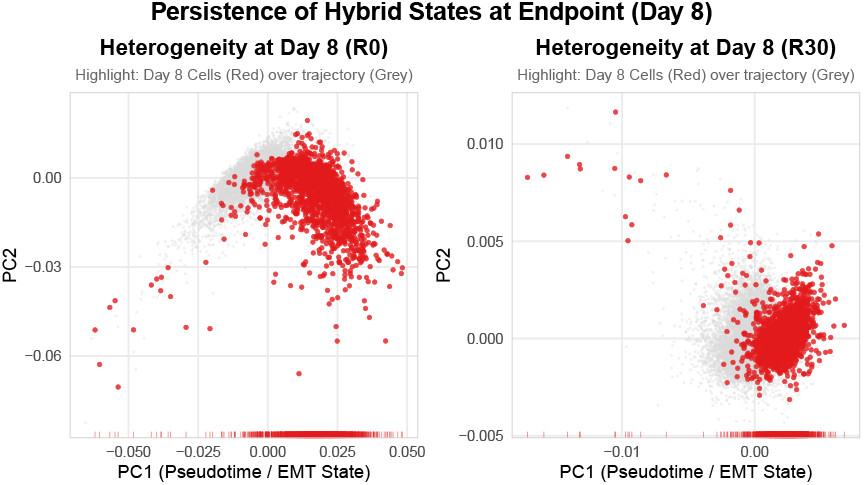
Cellular Heterogeneity at Day 8. Scatter plots showing that many Day 8 cells (red) remain in intermediate states along the trajectory, resisting complete transition.

## IV. Discussion

In this study, we proposed and validated an integrative computational pipeline that combines the network-based approach of *Transcriptogramer* with PCA dimensionality reduction. Beyond technical validation, the application of this method to the EMT model revealed that cellular plasticity is not an isolated cytoskeleton remodeling event but a **systemic reprogramming** coordinating bioenergetics, inflammatory response, and survival mechanisms.

### A. Network Topology as a Biological Noise Filter

A central challenge in scRNA-seq is the high dropout rate. Our results demonstrate that smoothing via the PPI network (R30 matrix) acts as a “biological low-pass filter.” By reconstructing a gene’s signal based on its functional neighborhood, we identified robust modules—such as Cluster 5 (Metal Response/Detoxification)—that would remain hidden in conventional gene-by-gene analyses. This validates the hypothesis that cellular functionality is an emergent property of network modules, not isolated genes.

### B. The Metabolic Switch: Powering Invasion

A major finding was the magnitude of metabolic reprogramming associated with EMT. While classical literature focuses on structural markers, our analysis showed that the most significant statistical enrichment (*p* < 10^*−*64^) occurred in **Cluster 2**, dominated by Oxidative Phosphorylation. This suggests a bioenergetic model where the EMT cell abandons the glycolytic profile (Warburg effect) and activates efficient mitochondrial respiration to sustain the high energy demand of motility.

### C. Mechanistic Validation: Cluster 5 and Oxidative Stress Response

The identification of Cluster 5, enriched for *Detoxification* and *Response to Metal Ion* (*MT1X, MT2A*), demonstrates our unsupervised pipeline’s ability to recover complex biological associations. Recent studies [8], [9] have linked heavy metals to EMT induction. Our results advance this understanding by showing that the EMT program itself intrinsically upregulates metal response genes. We propose a compensatory mechanism: the EMT cell activates a “dangerous power plant” (Cluster 2, mitochondrial respiration), which generates Reactive Oxygen Species (ROS), and simultaneously raises a “protective shield” (Cluster 5, metallothioneins) to scavenge these free radicals and prevent DNA damage. This molecular signature provides mechanistic support for the drug resistance often observed in metastatic cells.

## V. Conclusion

This study transcends the technical proposition of a new analysis method. By integrating *Transcriptogramer* with deep functional characterization, we revealed the modular architecture of the Epithelial-Mesenchymal Transition. We demonstrated that EMT is a systemic program coupling structural plasticity to a profound **bioenergetic reprogramming** and the intrinsic activation of **chemoresistance mechanisms**. The capacity of this method to automatically identify connections validated by recent literature confirms its robustness for knowledge discovery in sparse data. We conclude that the “functional map” generated by this approach offers a new translational perspective for oncology: effective targeting of metastasis may lie not only in blocking migration but in dismantling the metabolic supports and molecular defense shields that allow the cell to survive the transition.

## Supporting information

Full_Manuscript

## Data and code availability

The single-cell RNA-seq datasets analyzed in this study are publicly available in the Gene Expression Omnibus (GEO) under accession code GSE147405. The computational pipeline, including the Transcriptogramer analysis and figure generation scripts, is open-source and available at https://github.com/OdilonJulio/EMT_Transcriptogramer_Pipeline.

---

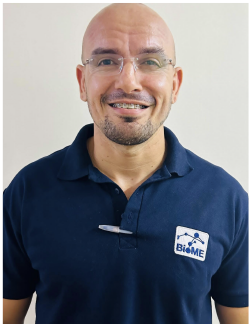

**Odilon Júlio dos Santos** received the B.S. degree in Mathematics from the Federal University of Rio Grande do Norte (UFRN), Brazil. He is currently working toward the M.Sc. degree with the Bioin-formatics Multidisciplinary Environment (BioME) at UFRN. His research interests include systems biology, network medicine, and computational analysis of single-cell transcriptomics.

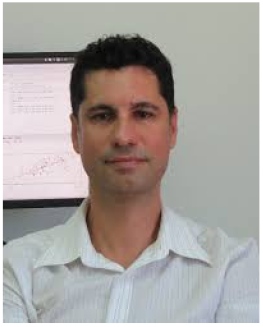

**Rodrigo J. S. Dalmolin** received the Ph.D. degree in Bioinformatics. He is currently an Associate Professor with the Bioinformatics Multidisciplinary Environment (BioME), UFRN. His research focuses on biological networks and systems biology.

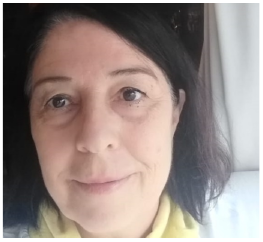

**Rita M. C. de Almeida** received the Ph.D. degree in Physics. She is a Full Professor at the Institute of Physics, UFRGS, and a collaborator at BioME/UFRN. Her research interests include statistical physics applied to biology and gene regulatory networks.

