## Supplementary material for "Analysis of Transcriptograms in Epithelial-Mesenchymal Transition (EMT)": Full_Manuscript: 00_main.tex

\documentclass[journal,twoside]{IEEEtran}

% --- Pacotes Oficiais IEEE & Essenciais ---
\usepackage{cite}
\usepackage{amsmath,amssymb,amsfonts}
\usepackage{algorithmic}
\usepackage{graphicx}
\usepackage{textcomp}
\usepackage{xcolor}
\usepackage{hyperref}
\usepackage{placeins}
\usepackage{float}
\usepackage[maxfloats=50]{morefloats} % Aumenta o limite

% --- Configuração de Subfiguras para IEEE ---
\usepackage[caption=false]{subfig}

% --- Caminho das Imagens ---
\graphicspath{{Figuras/}}

% --- Pacotes Gráficos e Ícones ---
\usepackage{tikz}
\usepackage{scalerel}
\usetikzlibrary{svg.path}
\usetikzlibrary{shapes.geometric, arrows, positioning, shadows.blur, calc}

% --- Definição do Ícone ORCID ---
\definecolor{orcidlogocol}{HTML}{A6CE39}
\tikzset{
  orcidlogo/.pic={
    \fill[orcidlogocol] svg{M256,128c0,70.7-57.3,128-128,128C57.3,256,0,198.7,0,128C0,57.3,57.3,0,128,0C198.7,0,256,57.3,256,128z};
    \fill[white] svg{M86.3,186.2H70.9V79.1h15.4v48.4V186.2z}
                 svg{M108.9,79.1h41.6c39.6,0,57,28.3,57,53.6c0,27.5-21.5,53.6-56.8,53.6h-41.8V79.1z M124.3,172.4h24.5c34.9,0,42.9-26.5,42.9-39.7c0-21.5-13.7-39.7-43.7-39.7h-23.7V172.4z}
                 svg{M88.7,56.8c0,5.5-4.5,10.1-10.1,10.1c-5.6,0-10.1-4.5-10.1-10.1c0-5.6,4.5-10.1,10.1-10.1C84.2,46.7,88.7,51.3,88.7,56.8z};
  }
}
\newbox{\orcidbox}
\sbox{\orcidbox}{\begin{tikzpicture}[yscale=-1,transform shape]\pic{orcidlogo};\end{tikzpicture}}
\newcommand\orcidicon[1]{\href{https://orcid.org/#1}{\mbox{\scalerel*{\usebox{\orcidbox}}{|}}}}

\begin{document}

% --- Título ---
\title{Analysis of Transcriptograms in Epithelial-Mesenchymal Transition (EMT)}

% --- Autores ---
\author{Odilon~Júlio~dos~Santos~\orcidicon{0009-0007-5081-3455},
        Rodrigo~Juliani~Siqueira~Dalmolin~\orcidicon{0000-0002-1688-6155},
        and~Rita~M.~C.~de~Almeida~\orcidicon{0000-0003-3897-1841}%
        
\thanks{Manuscript received \today; revised \today.}%
\thanks{The authors are with the Bioinformatics Multidisciplinary Environment (BioME), Federal University of Rio Grande do Norte (UFRN), Natal, Brazil.}%
}

\maketitle

%\markboth{IEEE Transactions on Computational Biology and Bioinformatics,~Vol.~XX, No.~X, February~2026}%
%{SANTOS \MakeLowercase{\textit{et al.}}: ANALYSIS OF TRANSCRIPTOGRAMS IN EPITHELIAL-MESENCHYMAL TRANSITION (EMT)}

% --- Corpo do Artigo ---
\input{01_abstract}
\input{02_introduction}
\input{03_methodology}
\input{04_results}
\input{05_discussion}

% --- Disponibilidade de Dados e Código (ATUALIZADO COM GITHUB) ---
\section*{Data and Code Availability}
The single-cell RNA-seq datasets analyzed in this study are publicly available in the Gene Expression Omnibus (GEO) under accession code GSE147405. The computational pipeline, including the Transcriptogramer analysis and figure generation scripts, is open-source and available at \url{https://github.com/OdilonJulio/EMT_Transcriptogramer_Pipeline}.

\FloatBarrier

% --- Referências ---
\bibliographystyle{IEEEtran}
\bibliography{references}

% --- Biografias ---
\begin{IEEEbiography}[{\includegraphics[width=1in,height=1.25in,clip,keepaspectratio]{odilon_foto.jpg}}]{Odilon Júlio dos Santos}
received the B.S. degree in Mathematics from the Federal University of Rio Grande do Norte (UFRN), Brazil. He is currently working toward the M.Sc. degree with the Bioinformatics Multidisciplinary Environment (BioME) at UFRN. His research interests include systems biology, network medicine, and computational analysis of single-cell transcriptomics.
\end{IEEEbiography}

\begin{IEEEbiography}[{\includegraphics[width=1in,height=1.25in,clip,keepaspectratio]{rodrigo_foto.png}}]{Rodrigo J. S. Dalmolin}
received the Ph.D. degree in Bioinformatics. He is currently an Associate Professor with the Bioinformatics Multidisciplinary Environment (BioME), UFRN. His research focuses on biological networks and systems biology.
\end{IEEEbiography}

\begin{IEEEbiography}[{\includegraphics[width=1in,height=1.25in,clip,keepaspectratio]{rita_foto.png}}]{Rita M. C. de Almeida}
received the Ph.D. degree in Physics. She is a Full Professor at the Institute of Physics, UFRGS, and a collaborator at BioME/UFRN. Her research interests include statistical physics applied to biology and gene regulatory networks.
\end{IEEEbiography}

\end{document}
