## Supplementary material for "Analysis of Transcriptograms in Epithelial-Mesenchymal Transition (EMT)": Full_Manuscript: 03_methodology.tex

\section{Methodology}

\subsection{Computational Pipeline Overview}
The analytical framework developed in this study combines network-based noise attenuation with dimensionality reduction to analyze single-cell transcriptomic data. As outlined in Fig. \ref{fig:pipeline_overview}, the workflow begins with raw data acquisition and stringent quality control, followed by application of the Transcriptogramer algorithm \cite{rybarczyk,morais}. The processed data are then subjected to PCA decomposition, enabling subsequent functional characterization of the resulting gene modules.

\begin{figure}[H]
    \centering
    \includegraphics[width=\linewidth]{pipeline_flowchart_new.png}
    \caption{\textbf{Integrative Computational Pipeline.} The flowchart details the sequential steps: (1) Pre-processing of raw scRNA-seq data; (2) The core \textit{Transcriptogramer} method acting as a biological low-pass filter; (3) PCA-based dimensionality reduction and trajectory reconstruction; (4) Functional clustering; and (5) The resulting biological insights regarding metabolism and inflammation.}
    \label{fig:pipeline_overview}
\end{figure}

\subsection{Data Acquisition and Batch Correction}
Publicly available scRNA-seq data from MCF10A cells treated with TGF-$\beta$1 were obtained from \cite{deshmukh}. The dataset comprises two batches of cells. The first one consists of samples treated up to 0, 4, and 8 days. The second batch consists of samples treated up to  0, 1, 2, and 3 days. The batches were processed in two different platforms (10x Genomics v2 and v3). Quality control was performed using the \texttt{Seurat} R package \cite{stuart2019}, filtering cells with fewer than 500 detected genes or mitochondrial gene content greater than 20\%.

To mitigate technical variability between the two experimental batches (Batch 1 and Batch 2), we applied a linear scaling correction based on the biological controls (Day 0 samples), which serve as a common baseline. Since both batches contain untreated cells representing the same biological state, systematic deviations in their mean gene expression profiles were attributed to batch effects.

For each gene $g$, we calculated the mean expression $\bar{X}^{(b)}_g$ for each batch $b$ using exclusively the Day 0 cells. We then defined a scaling factor $K_g$ as the ratio between the mean expression in the reference batch (Batch 2) and the target batch (Batch 1):

\begin{equation}
K_g = \frac{\bar{X}^{(2)}_g}{\bar{X}^{(1)}_g}
\end{equation}

The normalized expression values $X_{g,c}$ for all cells $c$ in Batch 1 were then adjusted as:

\begin{equation}
X^{\text{corr}}_{g,c} = X_{g,c} \times K_g
\end{equation}

This linear transformation aligns the basal expression levels of the controls across datasets while preserving the relative biological variation induced by the TGF-$\beta$1 treatment in subsequent time points. The correction was implemented in R using vectorized operations.

\subsection{The Transcriptogram Method}
The central component of our methodology is the Transcriptogramer package \cite{rybarczyk, morais}. In contrast to conventional approaches that consider genes as independent variables, this method projects gene expression data onto a gene list that has been ordered using information from Protein-Protein Interaction (PPI) networks (STRING-DB, score $\ge$ 800).

\subsubsection{Functional Ordering (CFM Algorithm)}
The linear ordering of genes is not arbitrary. It is obtained by minimizing a cost function $C$ that penalizes the distance between interacting proteins in the 1D list:
\begin{eqnarray}
C &=& \sum_{i<j} (j-i)^2 [ \, |A_{ij}-A_{ij-1}|+ |A_{ij}-A_{ij+1}| \nonumber \\
&+&|A_{ij}-A_{i-1j}|+|A_{ij}-A_{i+1j}| \, ] \,\, ,
\end{eqnarray}
where $A_{ij}$ is the adjacency matrix of the PPI network. This optimization is performed using a Monte Carlo algorithm (Simulated Annealing), ensuring that functionally related genes are placed in close proximity \cite{rybarczyk}.

\subsubsection{Biological Low-Pass Filter}
We applied a moving average window of radius $r=30$ (encompassing 61 genes) to smooth the expression profiles over this ordered list. This process acts as a \textbf{biological low-pass filter}:
\begin{equation}
t_g = \frac{1}{2r + 1}\sum_{i=g-r}^{g+r} E_i
\end{equation}
where $E_g$ is the expression of the gene at 

position $g$, where $t_g$ denotes the transcriptogram value corresponding to that gene. This smoothing procedure reduces stochastic noise (high-frequency fluctuations) inherent to scRNA-seq data while enhancing signals associated with coordinated functional modules (low-frequency variation) \cite{daSilva2014}.

Notably, setting $r=0$ yields transcriptograms in which raw expression values are assigned to each gene, without any smoothing. Even in this case, however, the result is not identical to the original expression matrix, since we restrict the analysis to genes included in the ordered list—namely, those whose protein products are recorded in the STRING database as having at least one protein–protein interaction with a score of 800 or higher. This preliminary filtering step reduces the number of genes from approximately 30,000 to about 14,000.

\subsection{PCA and Trajectory Reconstruction}
Principal Component Analysis (PCA) was applied to the smoothed transcriptograms to achieve dimensionality reduction. Transcriptomes were then reconstructed using a subset of principal components (PCs), and the reconstruction error was evaluated by comparing the reconstructed transcriptograms with the original ones. The expression for reconstructing the transcriptome of a cell $c$ using the first 
$k$ components is given by:
\begin{equation}
\mathbf{t}_{rec}^{(c)} = \boldsymbol{\mu} + \sum_{i=1}^{k} (S_{c,i} \cdot \mathbf{v}_i)
\end{equation}
where $\boldsymbol{\mu}$ and $\mathbf{t}_{rec}^{(c)}$ are, respectively, the mean expression vector and the reconstructed transcriptogram vector,  $S_{c,i}$ is the coefficiet of cell $c$ in the durection defined by  PC $i$, and $\mathbf{v}_i$ is the $i$th principal component. This step allows for the visualization of global "waves" of gene expression along the EMT progression.

\subsection{Functional Characterization and "Rescue" Pipeline}
To interpret the biological patterns, we identified regions of high variance between samples  along the transcriptograms. We then clustered them into functional modules using k-means. These clusters were initially characterized using the MSigDB Hallmark collection via the \texttt{clusterProfiler} package.

To overcome the limited coverage of general Hallmarks, we implemented a \textbf{"Rescue Pipeline"}. For clusters showing no significant Hallmark enrichment (e.g., Cluster 5), we automatically queried the Gene Ontology (GO) Biological Processes database. This hierarchical approach allowed us to identify specific mechanisms, such as "Response to Metal Ion" and "Detoxification", which are critical for cell survival but often overlooked in EMT standard broad-spectrum analyses.
