## Supplementary material for "Analysis of Transcriptograms in Epithelial-Mesenchymal Transition (EMT)": Full_Manuscript: 04_results.tex

\section{Results}

\subsection{Method Validation: Dimensionality Reduction Efficiency}
Before dissecting the biological mechanisms, we evaluated the capacity of the Transcriptogramer method using $r=30$ (R30 samples) to compress relevant biological information. 

However, the smoothing step is decisive for signal consolidation. As shown in \textbf{Fig. \ref{fig:pca_variance_r30}}, the variance decomposition analysis of the R30 transcriptograms reveals a rapid accumulation of the explained variance. The detailed view of the first 25 components \textbf{(Left Panel)} clearly identifies the "elbow point" (saturation of the relevant topological signal), demonstrating that the core biological trajectory is captured within the first few dimensions. The global spectrum \textbf{(Right Panel)} confirms that higher-order components contribute marginally, mostly representing stochastic noise. This efficiency justifies the use of a low-dimensional space for the trajectory reconstruction in the subsequent analyses.

\begin{figure*}[!ht]
    \centering
    \includegraphics[width=0.95\textwidth]{pca_variance_R30_focused_25PC.pdf}
    \caption{\textbf{Structural Decomposition of Variance in R30 Condition.} 
    \textbf{(Left)} Detailed analysis of the first 25 Principal Components highlights the "elbow point," indicating where the accumulation of biological signal saturates. 
    \textbf{(Right)} The global spectrum (all components) shows the long tail of low-variance dimensions, confirming that topological smoothing effectively compresses the system's dynamics into the leading components, filtering out noise.}
    \label{fig:pca_variance_r30}
\end{figure*}

\subsection{Trajectory Reconstruction}
Fig. \ref{fig:pca_multipanel_r30} presents the plots of the values of coefficients of several principal components as a function of the coefficients of the first principal component, PC1.
\begin{figure*}[!ht]
    \centering
    \includegraphics[width=0.95\textwidth]{Figuras/pca_grid_PC1_vs_PC2to6_R30.pdf}
    
    \caption{\textbf{Structural Decomposition of Variance and Trajectory Analysis (R30).}     Plots of higher-order components coefficients (PC2 to PC6) against first Principal Component coefficient (PC1). The black arrows connect the centroids of each time point, illustrating the trajectory of the system's center of mass from Day 0 to Day 8.
    \textbf{(A-C, Top Row)} PC2, PC3, and PC4 vs PC1 reveal a robust temporal structure, with clear separation between early (purple/blue), intermediate (green), and late (orange/red) stages.
    \textbf{(D-E, Bottom Row)} As dimensionality increases (PC5 and PC6), the temporal segregation decreases, confirming that the main biological signal driving the EMT is concentrated in the first few dimensions. The color legend (bottom right) applies to all panels.}
    \label{fig:pca_multipanel_r30}
\end{figure*}

\subsection{Global Dynamics: Waves of Expression}
We first plot in Fig. 2 the values of coeficients relative to different components (PC2 up to  PC6)  as a function of the coeficient for the first principal component PC1 for  all the samples, in different collors for each day. As shown in Fig. 1, upper, left panel, components 1 to 6 concentrate more than 75\% of the total variance. There is a clear progression from smaller to larger values of PC1 coeficient for samples harvested at later days, showing that we can use this parameter as a proxy for EMT progression.  The resulting analysis reveals that EMT does not occur as an abrupt switch, but as a pattern of "expression waves" as the cells progress in EMT (Fig. \ref{fig:transcriptogram_storyboard}). This is useful for further analyses.

To visualize the continuous nature of the transition, we reconstructed all transcriptomes using 6 principal components (from PC1 to PC6). Then, we subdivided the samples by their PC1 values into 50 intervals.. For the samples localized at the same PC1 coefficient interval, we calculated the average transcriptograms. This is shown in Fig \textbf{Fig. \ref{fig:transcriptogram_storyboard}}, for different stages of EMT progression. The transcriptograms in this figure represents the expression changes in respect to the global transcriptogram average, or the center of mass of the points representing the samples in the expression space used in PCA. Hence, zero value in these transcriptograms represents zero change in respect to the average sample. As background, these figures show pathways or GO terms localization on the gene ordering. This is meant to help the biological interpretation of the observed gene expression alterations. 

\begin{figure*}[!htbp]
    \centering
    
    % --- Linha 1 ---
    \subfloat{\includegraphics[width=0.5\linewidth]{Fig_Part_A.png}}\hfill
    \subfloat{\includegraphics[width=0.5\linewidth]{Fig_Part_B.png}} \\
    
    % --- Linha 2 ---
    \subfloat{\includegraphics[width=0.5\linewidth]{Fig_Part_C.png}}\hfill
    \subfloat{\includegraphics[width=0.5\linewidth]{Fig_Part_D.png}} \\

    % --- Linha 3 ---
    \subfloat{\includegraphics[width=0.5\linewidth]{Fig_Part_E.png}}\hfill
    \subfloat{\includegraphics[width=0.5\linewidth]{Fig_Part_F.png}} \\

    % --- Linha 4 ---
    \subfloat{\includegraphics[width=0.5\linewidth]{Fig_Part_G.png}}\hfill
    \subfloat{\includegraphics[width=0.5\linewidth]{Fig_Part_H.png}} \\
    
    % --- LEGENDA --
    
    \vspace{0.5em} 
    
    \includegraphics[width=0.8\linewidth]{Figuras/Fig_Legend.png} 
    
    \caption{\textbf{Evolutionary Panel of Global Transcriptome Reconstruction.} The sequence of panels illustrates the topological dynamics of gene expression waves along the ordered gene list. Valley regions (low expression) in early phases progressively transform into peaks (high expression) in later phases. The color bar (bottom) indicates the relative expression levels. \textbf{(A)} 0\% - Epithelial State; \textbf{(B)} 14\% - Initiation; \textbf{(C)} 28\% - Early Transition; \textbf{(D)} 42\% - Plasticity Peak I; \textbf{(E)} 57\% - Plasticity Peak II; \textbf{(F)} 71\% - Late Transition; \textbf{(G)} 85\% - Mesenchymal Est.; \textbf{(H)} 100\% - Mesenchymal Final.}
    \label{fig:transcriptogram_storyboard}
\end{figure*}

\subsection{Dynamic Orchestration of Functional Modules}
The gene variance detection algorithm segmented the transcriptogram into co-regulated gene clusters. To understand the temporal hierarchy of these modules, we analyzed their relative contribution to the global variance along the EMT progression.

Fig. \ref{fig:cluster_dynamics} illustrates this "changing of the guard". We observe a sequential orchestration where modules responsible for epithelial maintenance decline, giving way to modules driving the mesenchymal phenotype. Crucially, the normalized importance plot (Fig. \ref{fig:cluster_dynamics} TOP) and the stacked area plot (Fig. \ref{fig:cluster_dynamics} BOTTOM) reveal that the transition is supported by a massive metabolic shift (Cluster 2) that persists and expands towards the end of the trajectory.

\begin{figure}[!htbp]
    \centering
    % Imagem Superior
    \subfloat{\includegraphics[width=\linewidth]{importancia_normalizada_por_frame.png}}\\
    % Imagem Inferior
    \subfloat{\includegraphics[width=\linewidth]{area_stacked_clusters.png}}
    \caption{\textbf{Dynamics of Functional Modules.} 
    \textbf{(Top)} Normalized Relative Importance plot showing the shifting dominance of specific gene clusters along the pseudotime. 
    \textbf{(Bottom)} Stacked Contribution Area plot confirming that EMT is a continuous replacement of genetic programs, with no periods of transcriptional silence. Note the expansion of Cluster 2 (Metabolism) and Cluster 6 (Core EMT/Inflammation) in the later stages.}
    \label{fig:cluster_dynamics}
\end{figure}

\subsection{Characterization of Key Modules}

\subsubsection{The "Core" EMT and Inflammation (Cluster 6)}
Cluster 6 was identified as the central driver, showing robust enrichment for the \textit{Epithelial Mesenchymal Transition} signature ($p < 10^{-19}$) and inflammatory pathways (\textit{TNFA signaling via NFKB}), suggesting inflammation is intrinsic to the mesenchymal state (Fig. \ref{fig:panel_final}).

\begin{figure*}[!htbp]  
    \centering
    \includegraphics[width=\textwidth]{PANEL_FINAL_ARTIGO.png}   
    \caption{\textbf{Functional Architecture of EMT.} (A) Statistical validation of EMT signature in Clusters 3 and 6. (B) Global landscape revealing parallel processes. (C) Heatmap of the functional pillars: Plasticity, Bioenergetics, Inflammation, and Stress.}
    \label{fig:panel_final}
\end{figure*}

\subsubsection{The Metabolic Switch (Cluster 2)}
Cluster 2 represents a purely metabolic module, massively enriched for \textit{Oxidative Phosphorylation} ($p < 10^{-64}$). This confirms the bioenergetic reprogramming required to sustain the high energy demands of invasion.

\subsubsection{The Detoxification Shield (Cluster 5)}
Cluster 5 was rescued by our Gene Ontology pipeline, revealing functions related to \textit{Response to Metal Ion} and \textit{Detoxification}. The upregulation of metallothioneins (\textit{MT1X}, \textit{MT2A}) indicates an endogenous chemoresistance mechanism activated during the transition.


\subsection{Comparative Analysis and Population Heterogeneity}
We compared our findings with canonical markers described by Deshmukh et al. \cite{deshmukh}. As shown in Fig. \ref{fig:venn_comparativo}, while there is convergence on structural markers, our network-based approach uniquely highlights the Metabolic Switch and Defense Shield.

\begin{figure}[H]
    \centering
    \includegraphics[width=\linewidth]{venn_comparativo.png}
    \caption{\textbf{Comparative Analysis of Gene Scope: Transcriptogramer vs. Literature.} 
    Overlap analysis between the topological clusters identified in this study and the classical EMT markers (Deshmukh et al., 2021). 
    (1) \textbf{Consensus:} Core EMT effectors were successfully retrieved, confirming method robustness. 
    (2) \textbf{Novel Insights:} Our approach uniquely prioritized mitochondrial (Cluster 2) and stress-response genes (Cluster 5), suggesting that metabolic reprogramming and oxidative stress management are integral components of the EMT program. 
    (3) \textbf{Literature-Exclusive:} Markers such as \textit{CDH1} (epithelial) were excluded by the topological filter as they represent the downregulated state rather than the active mesenchymal transition.}
    \label{fig:venn_comparativo}
\end{figure}


Finally, analysis at the experimental endpoint (Day 8) revealed marked heterogeneity (Fig. \ref{fig:day8_heterogeneity}). A significant fraction of cells did not reach the mesenchymal extreme, confirming the existence of stable hybrid phenotypes. This persistent hybrid population creates a reservoir of plasticity potentially linked to tumor dormancy or relapse.

\begin{figure}[H]
    \centering
    \includegraphics[width=\linewidth]{day8_heterogeneity_analysis.pdf}
    \caption{\textbf{Cellular Heterogeneity at Day 8.} Scatter plots showing that many Day 8 cells (red) remain in intermediate states along the trajectory, resisting complete transition.}
    \label{fig:day8_heterogeneity}
\end{figure}
