## Supplementary material for "Analysis of Transcriptograms in Epithelial-Mesenchymal Transition (EMT)": Full_Manuscript: references.bib

@article{dongre,
  author    = {Dongre, A. and Weinberg, R. A.},
  title     = {New insights into the mechanisms of epithelial–mesenchymal transition and implications for cancer},
  journal   = {Nature Reviews Molecular Cell Biology},
  volume    = {20},
  pages     = {69--84},
  year      = {2019},
  doi       = {10.1038/s41580-018-0080-4}
}

@article{morais,
  author    = {Morais, D. A. A. and Almeida, R. M. C. and Dalmolin, R. J. S.},
  title     = {Transcriptogramer: an R/Bioconductor package for transcriptional analysis based on protein–protein interaction},
  journal   = {Bioinformatics},
  volume    = {35},
  number    = {16},
  pages     = {2875--2876},
  year      = {2019},
  doi       = {10.1093/bioinformatics/btz004}
}

@article{rybarczyk,
  author    = {Rybarczyk-Filho, J. L. and Castro, M. A. and Dalmolin, R. J. S. and Moreira, J. C. and Brunnet, L. G. and de Almeida, R. M.},
  title     = {Towards a genome-wide transcriptogram: the Saccharomyces cerevisiae case},
  journal   = {Nucleic Acids Research},
  volume    = {39},
  number    = {8},
  pages     = {3005--3016},
  year      = {2011},
  doi       = {10.1093/nar/gkq1266}
}

@article{deshmukh,
  author    = {Deshmukh, A. P. and Vasaikar, S. V. and Tomczak, K. and Tripathi, S. and Den Hollander, P. and Arslan, E. and Chakraborty, P. and Soundararajan, R. and Jolly, M. K. and Rai, K. and Levine, H. and Mani, S. A.},
  title     = {Identification of EMT signaling cross-talk and gene regulatory networks by single-cell RNA sequencing},
  journal   = {Proceedings of the National Academy of Sciences},
  volume    = {118},
  number    = {19},
  pages     = {e2102050118},
  year      = {2021},
  doi       = {10.1073/pnas.2102050118}
}

@article{fontclos,
  author    = {Font-Clos, F. and Zapperi, S. and La Porta, C. A. M.},
  title     = {Topography of Epithelial-Mesenchymal Plasticity},
  journal   = {Frontiers in Bioengineering and Biotechnology},
  volume    = {8},
  pages     = {220},
  year      = {2020},
  doi       = {10.3389/fbioe.2020.00220}
}

@article{hari,
  author    = {Hari, K. and Sabhachandani, P. and Jolly, M. K.},
  title     = {Identifying inhibitors of epithelial–mesenchymal plasticity using a network topology-based approach},
  journal   = {Proceedings of the National Academy of Sciences},
  volume    = {119},
  number    = {26},
  pages     = {e2216109120},
  year      = {2022},
  doi       = {10.1073/pnas.2216109120}
}

@incollection{deAlmeida2020,
  author    = {De Almeida, R. M. C. and De Souza, L. L. S. and Morais, D. and Dalmolin, R. J. S.},
  title     = {Transcriptograms: A Genome-Wide Gene Expression Analysis Method},
  booktitle = {Networks in Systems Biology},
  editor    = {Da Silva, F. A. B. and Carels, N. and Trindade dos Santos, M. and Lopes, F. J. P.},
  series    = {Computational Biology},
  volume    = {32},
  publisher = {Springer, Cham},
  pages     = {117--132},
  year      = {2020},
  doi       = {10.1007/978-3-030-51862-2_5}
}

@manual{RCoreTeam2023,
  title        = {R: A Language and Environment for Statistical Computing},
  author       = {{R Core Team}},
  organization = {R Foundation for Statistical Computing},
  address      = {Vienna, Austria},
  year         = {2023},
  url          = {https://www.R-project.org/}
}

@manual{RStudio2023,
  title        = {RStudio: Integrated Development Environment for R},
  author       = {{RStudio Team}},
  year         = {2023},
  url          = {https://www.rstudio.com/}
}

@article{stuart2019,
  author  = {Stuart, T. and Butler, A. and Hoffman, P. and Hafemeister, C. and Papalexi, E. and Mauck, W. M. and Hao, Y. and Stoeckius, M. and Smibert, P. and Satija, R.},
  title   = {Comprehensive Integration of Single-Cell Data},
  journal = {Cell},
  volume  = {177},
  number  = {7},
  pages   = {1888--1902},
  year    = {2019},
  doi     = {10.1016/j.cell.2019.05.031}
}

@article{germain2022,
  author  = {Germain, P.-L. and Lun, A. T. L. and Macnair, W. and Robinson, M. D.},
  title   = {Doublet identification in single-cell sequencing data using scDblFinder},
  journal = {Nature Methods},
  volume  = {19},
  pages   = {279--281},
  year    = {2022}
}

@article{durinck2009,
  author  = {Durinck, S. and Spellman, P. T. and Birney, E. and Huber, W.},
  title   = {Mapping identifiers for the integration of genomic datasets with the biomaRt Bioconductor package},
  journal = {BMC Bioinformatics},
  volume  = {10},
  number  = {Suppl 1},
  pages   = {S4},
  year    = {2009},
  doi     = {10.1186/1471-2105-10-S1-S4}
}

@book{wickham2016,
  author    = {Wickham, H.},
  title     = {ggplot2: Elegant Graphics for Data Analysis},
  publisher = {Springer-Verlag New York},
  year      = {2016},
  isbn      = {978-3-319-24277-4}
}

@manual{NPAD2025,
  title        = {Núcleo de Processamento de Alto Desempenho (NPAD/UFRN)},
  author       = {{NPAD/UFRN}},
  organization = {Universidade Federal do Rio Grande do Norte},
  year         = {2025},
  url          = {https://npad.ufrn.br/npad/bemvindo},
  note         = {Acesso em: 2 ago. 2025}
}

@article{jolliffe2016,
  author    = {Jolliffe, I. T. and Cadima, J.},
  title     = {Principal Component Analysis: A Review and Recent Developments},
  journal   = {Philosophical Transactions of the Royal Society A},
  year      = {2016},
  volume    = {374},
  number    = {2065},
  pages     = {20150202}
}

@article{hotelling1933,
  author  = {Hotelling, H.},
  title   = {Analysis of a complex of statistical variables into principal components},
  journal = {Journal of Educational Psychology},
  volume  = {24},
  number  = {6},
  pages   = {417--441},
  year    = {1933}
}

@manual{Rparallel,
  title        = {parallel: Support for Parallel Computation in R},
  author       = {{R Core Team}},
  organization = {R Foundation for Statistical Computing},
  address      = {Vienna, Austria},
  year         = {2023},
  note         = {Part of R 4.3.1}
}

@article{yu2012,
  title={clusterProfiler: an R package for comparing biological themes among gene clusters},
  author={Yu, Guangchuang and Wang, Li-Gen and Han, Yanyan and He, Qing-Yu},
  journal={Omics: a journal of integrative biology},
  volume={16},
  number={5},
  pages={284--287},
  year={2012},
  publisher={Mary Ann Liebert, Inc. 140 Huguenot Street, 3rd Floor New Rochelle, NY 10801 USA}
}

@article{liberzon2015,
  title={The molecular signatures database (MSigDB) hallmark gene set collection},
  author={Liberzon, Arthur and Birger, Chet and Thorvaldsd{\'o}ttir, Helga and Ghandi, Mahmoud and Mesirov, Jill P and Tamayo, Pablo},
  journal={Cell systems},
  volume={1},
  number={6},
  pages={417--425},
  year={2015},
  publisher={Elsevier}
}

@article{ashburner2000,
  title={Gene ontology: tool for the unification of biology},
  author={Ashburner, Michael and Ball, Catherine A and Blake, Judith A and Botstein, David and Butler, Heather and Cherry, J Michael and Davis, Allan P and Dolinski, Kara and Dwight, Selina S and Eppig, Janan T and others},
  journal={Nature genetics},
  volume={25},
  number={1},
  pages={25--29},
  year={2000},
  publisher={Nature Publishing Group}
}

@article{carlson2019,
  title={org.Hs.eg.db: Genome wide annotation for Human},
  author={Carlson, Marc},
  journal={R package version},
  volume={3},
  number={8},
  year={2019}
}

@article{chen2022,
  title={Toxic metals in the regulation of epithelial--mesenchymal plasticity: demons or angels?},
  author={Chen, X-L and Xu, Y-M and Lau, ATY},
  journal={Cancer Cell International},
  volume={22},
  number={1},
  pages={1--15},
  year={2022},
  publisher={Springer}
}

@article{valverde2024,
  title={Epithelial--Mesenchymal Transition Induced by a Metal Mixture in Liver Cells (As, Cd, Pb)},
  author={Valverde, M and Rosales-Cruz, P and others},
  journal={Oxidative Medicine and Cellular Longevity},
  volume={2024},
  year={2024},
  publisher={Hindawi}
}

@article{chang2019,
  title={Arsenic-induced neoplastic transformation involves EMT and $\beta$-catenin/c-Myc pathway},
  author={Chang, Y-W and others},
  journal={Chemical Research in Toxicology},
  volume={32},
  number={5},
  pages={849--859},
  year={2019},
  publisher={ACS Publications}
}

@article{shi2021,
  title={Cadmium induces EMT in renal cancer cells via PGE2/cAMP signaling},
  author={Shi, H and Sun, X and others},
  journal={Ecotoxicology and Environmental Safety},
  volume={208},
  pages={111608},
  year={2021},
  publisher={Elsevier}
}

@article{daSilva2014,
  author  = {da Silva, S. R. M. and Perrone, G. C. and Dinis, J. M. and de Almeida, R. M. C.},
  title   = {Transcriptograms: reproducibility enhancement and differential expression of non-predefined functional gene sets in human genome},
  journal = {BMC Genomics},
  volume  = {15},
  number  = {1},
  pages   = {1181},
  year    = {2014},
  doi     = {10.1186/1471-2164-15-1181}
}
