## Supplementary figures and images for "Analysis of Transcriptograms in Epithelial-Mesenchymal Transition (EMT)"

### area_stacked_clusters.png

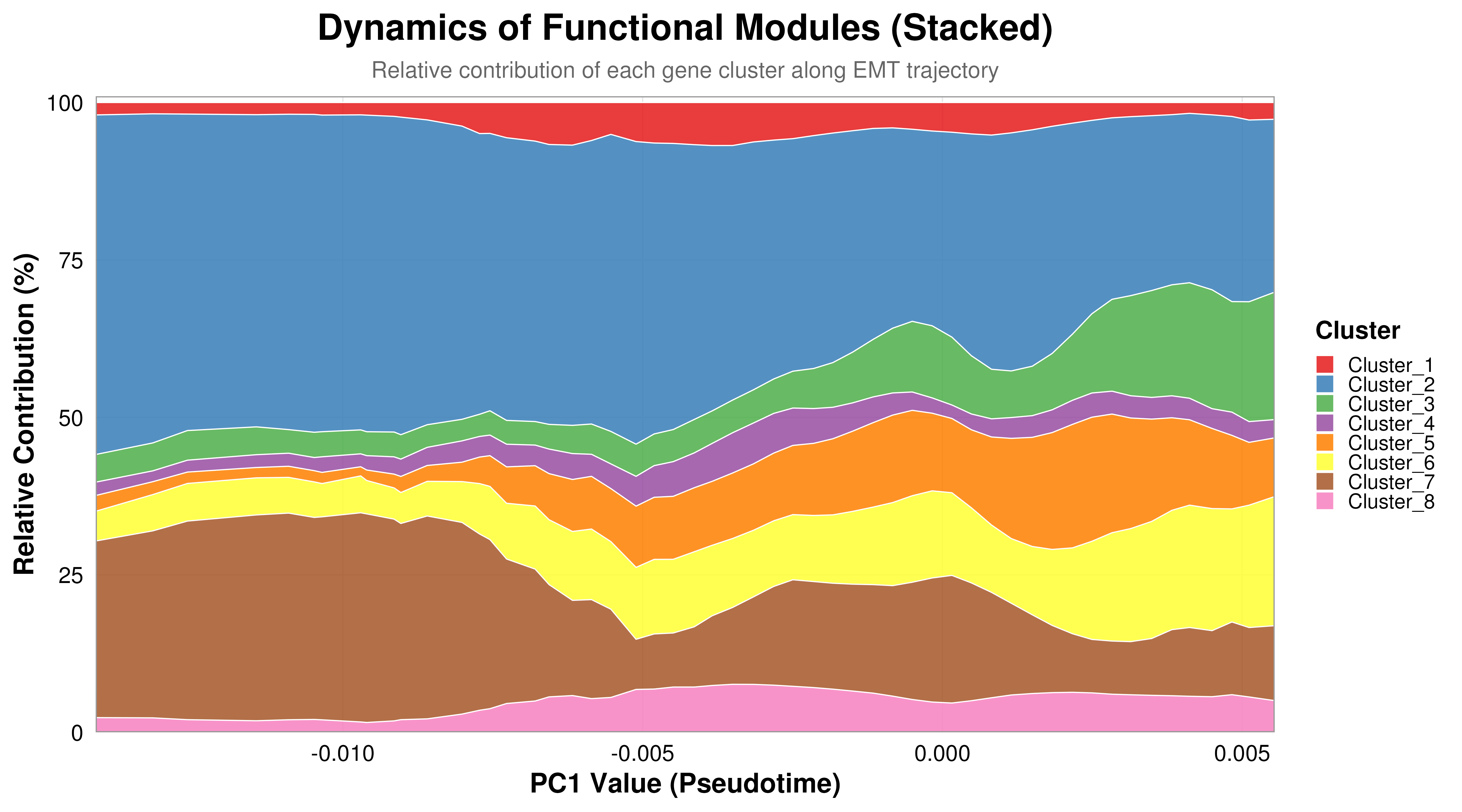

### day8_heterogeneity_analysis.pdf

#### Persistence of Hybrid States at Endpoint (Day 8)

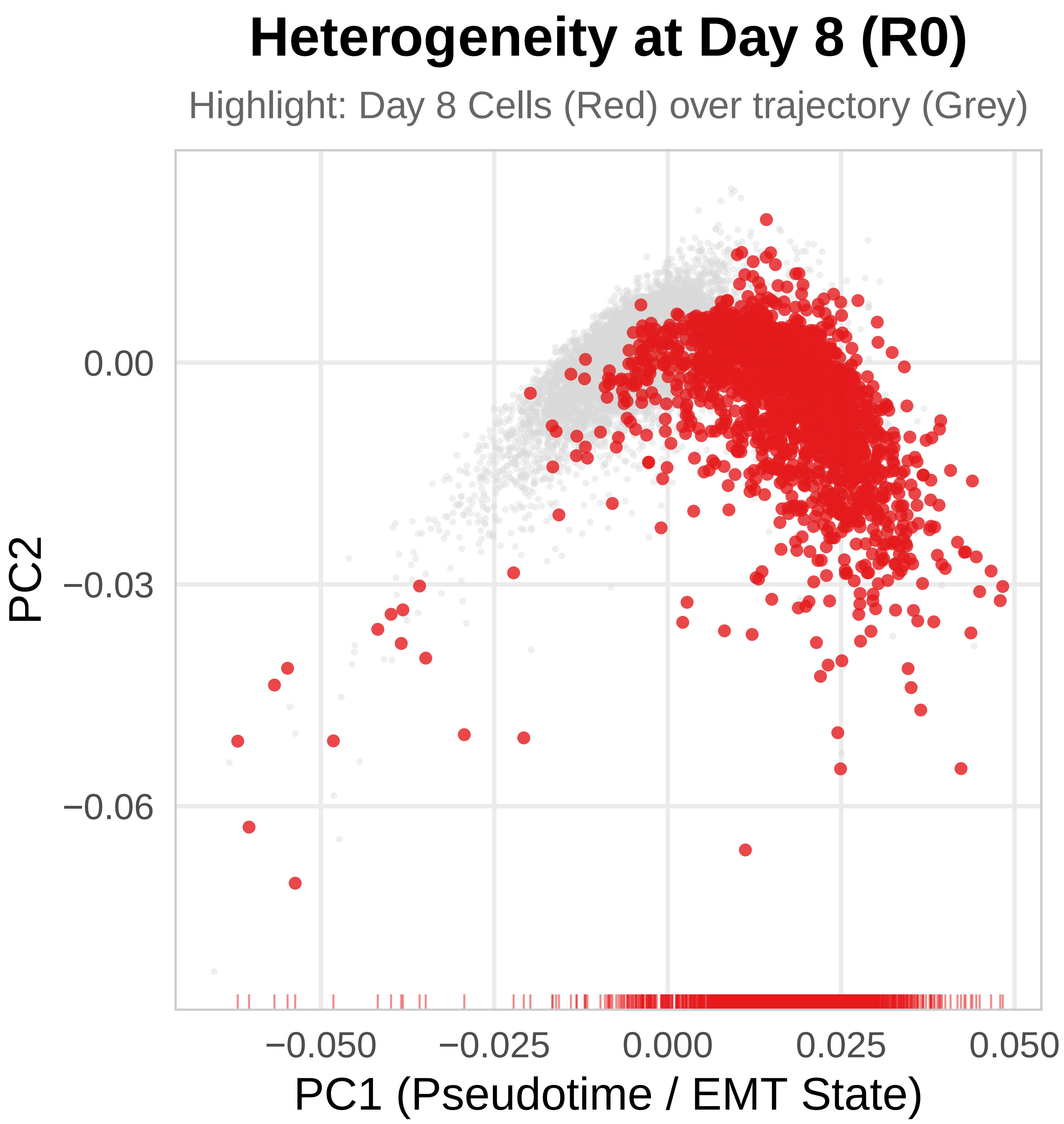


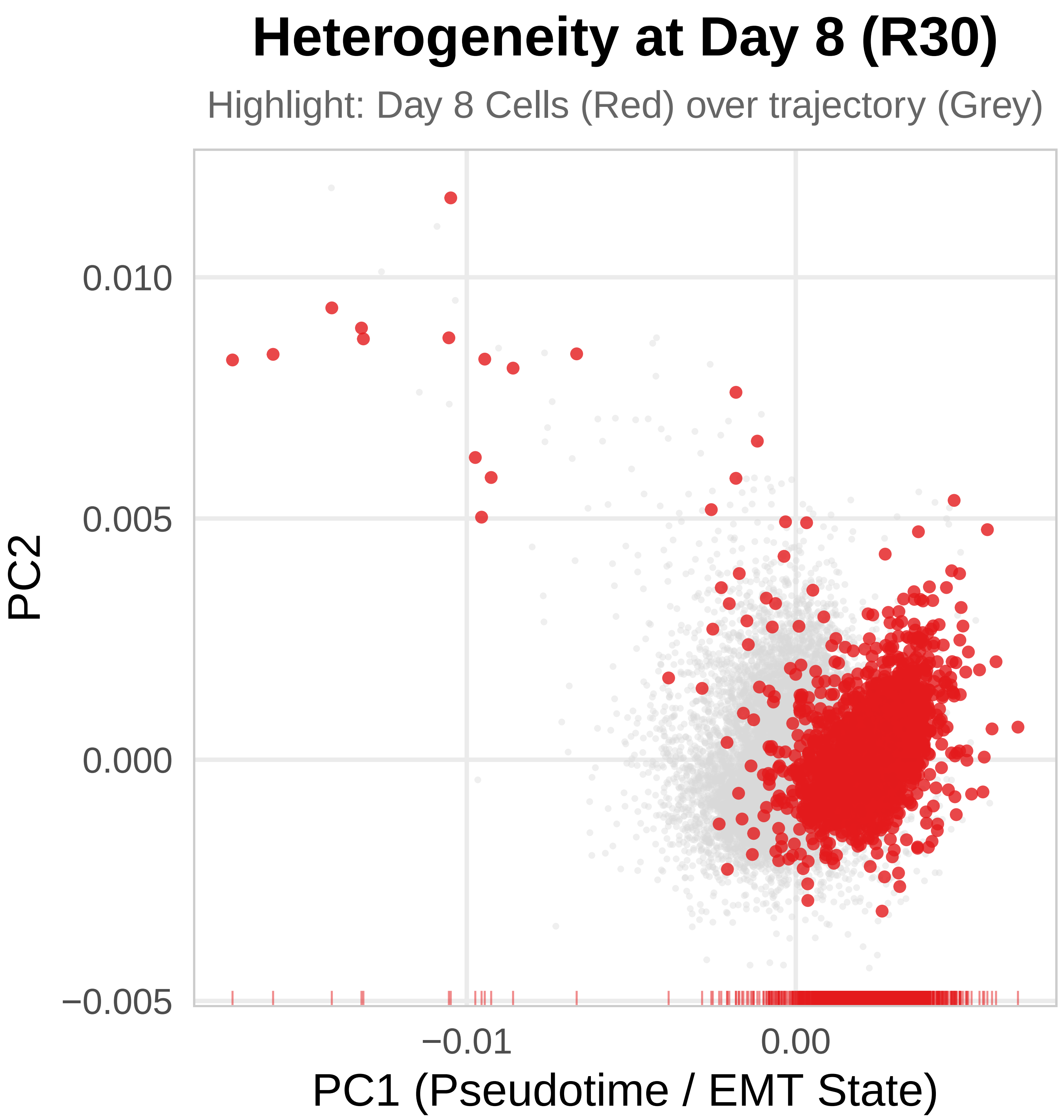

### Fig_Legend.png

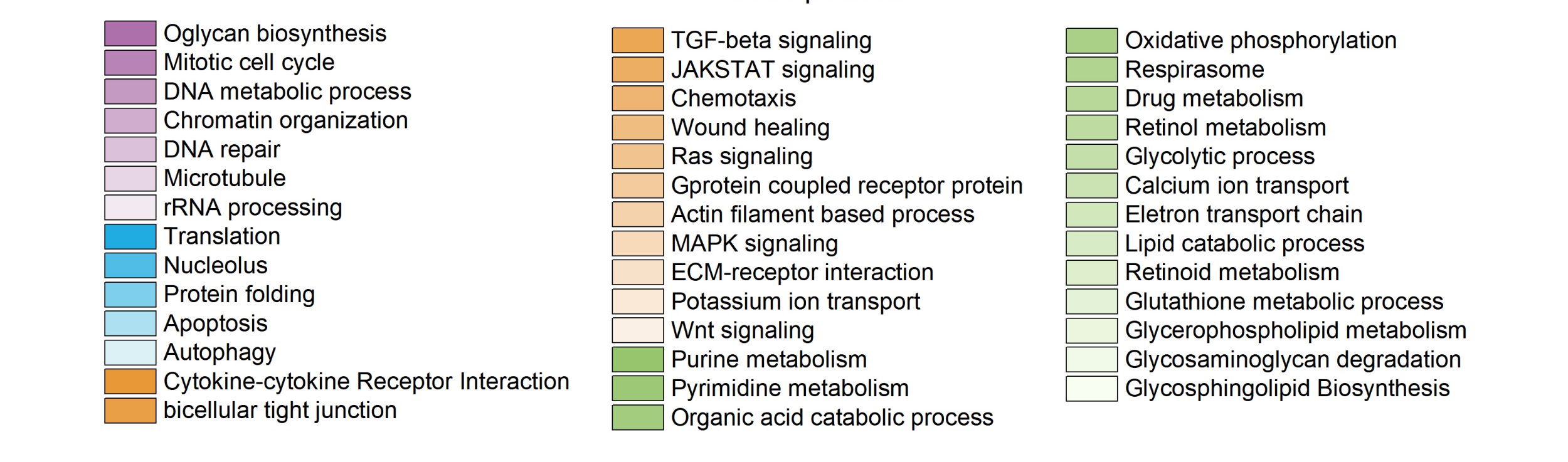

### Fig_Part_A.png

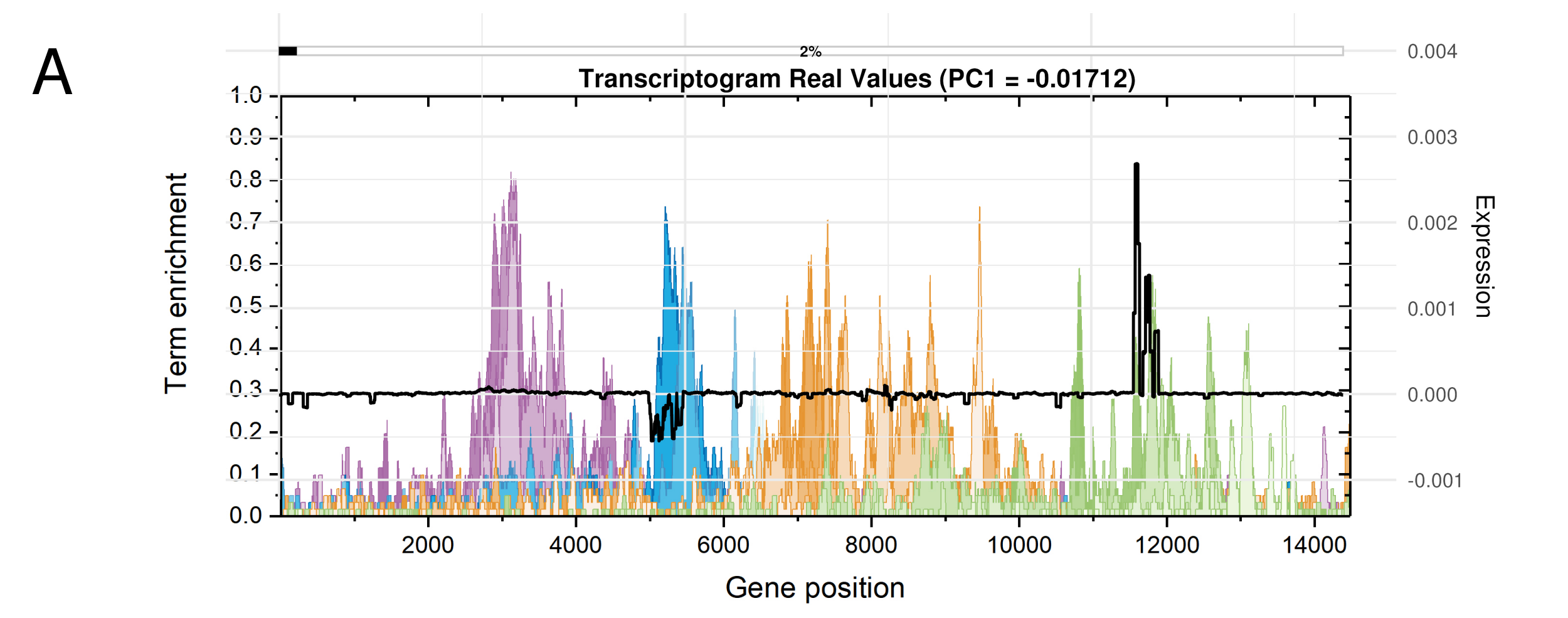

### Fig_Part_B.png

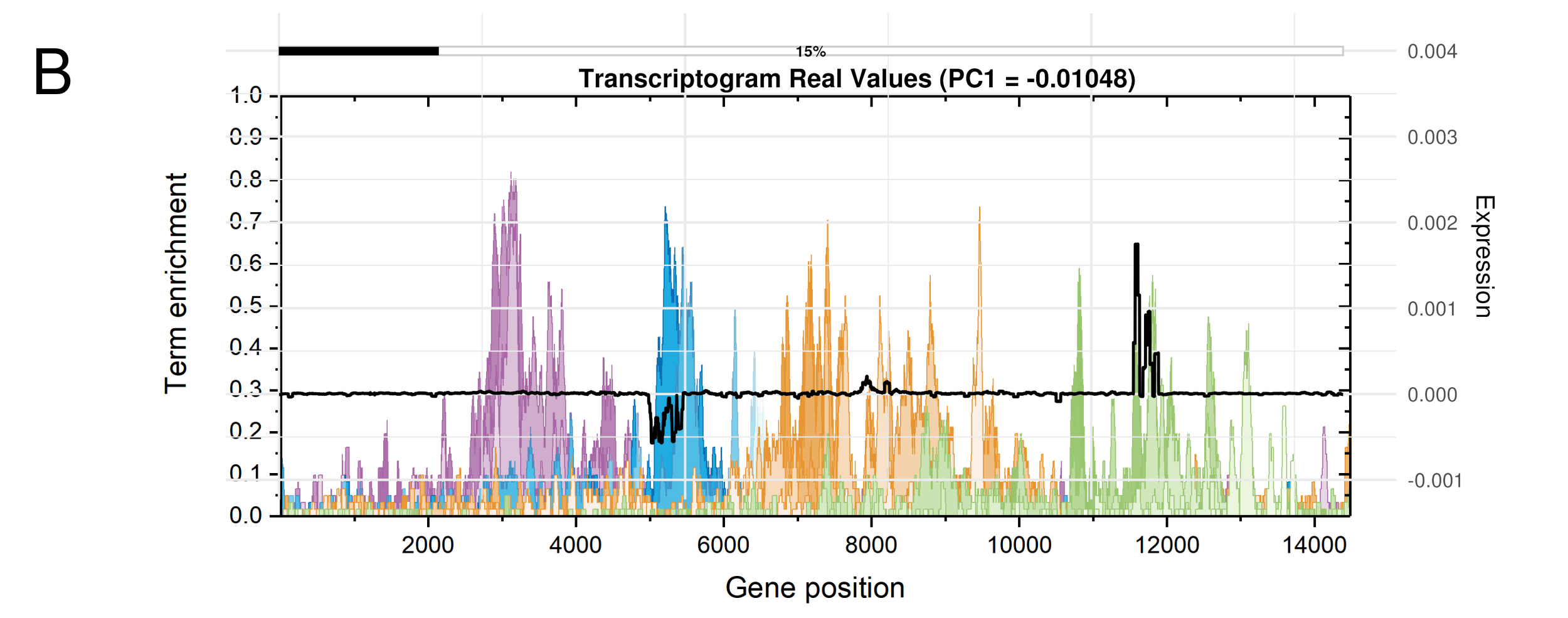

### Fig_Part_C.png

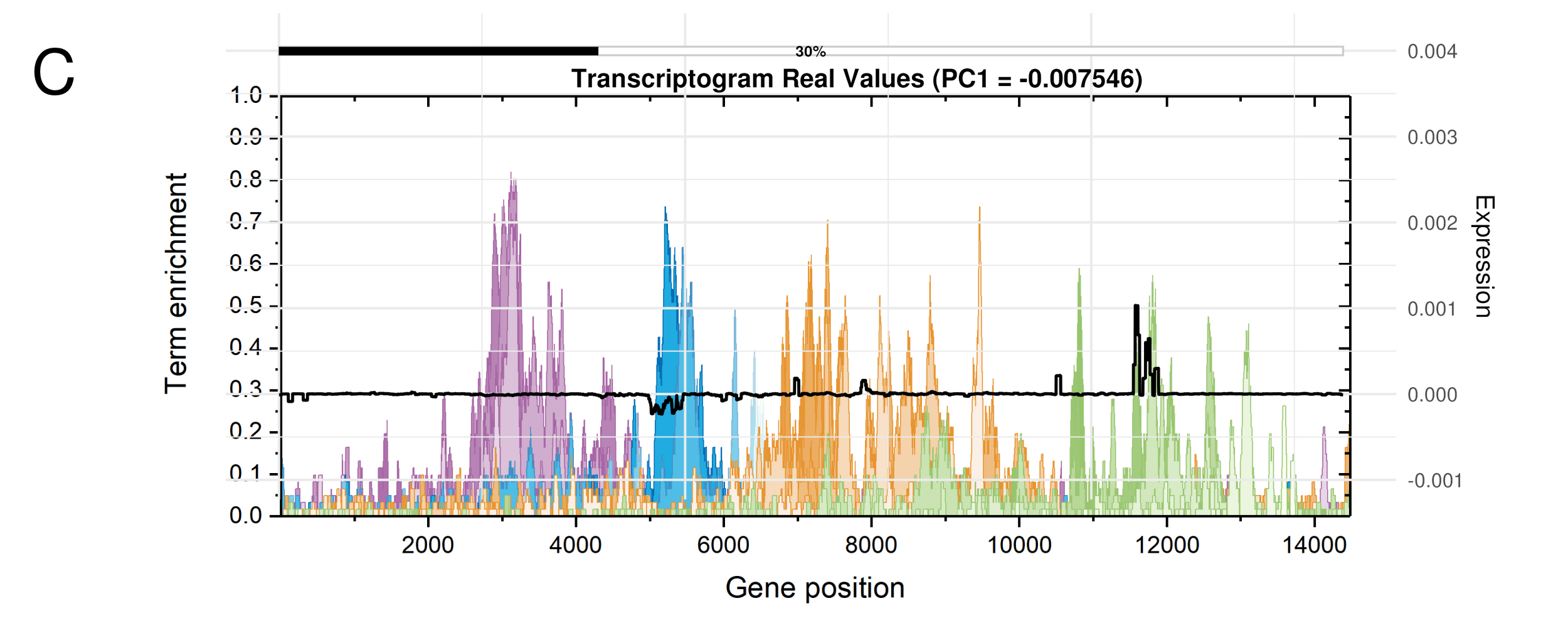

### Fig_Part_D.png

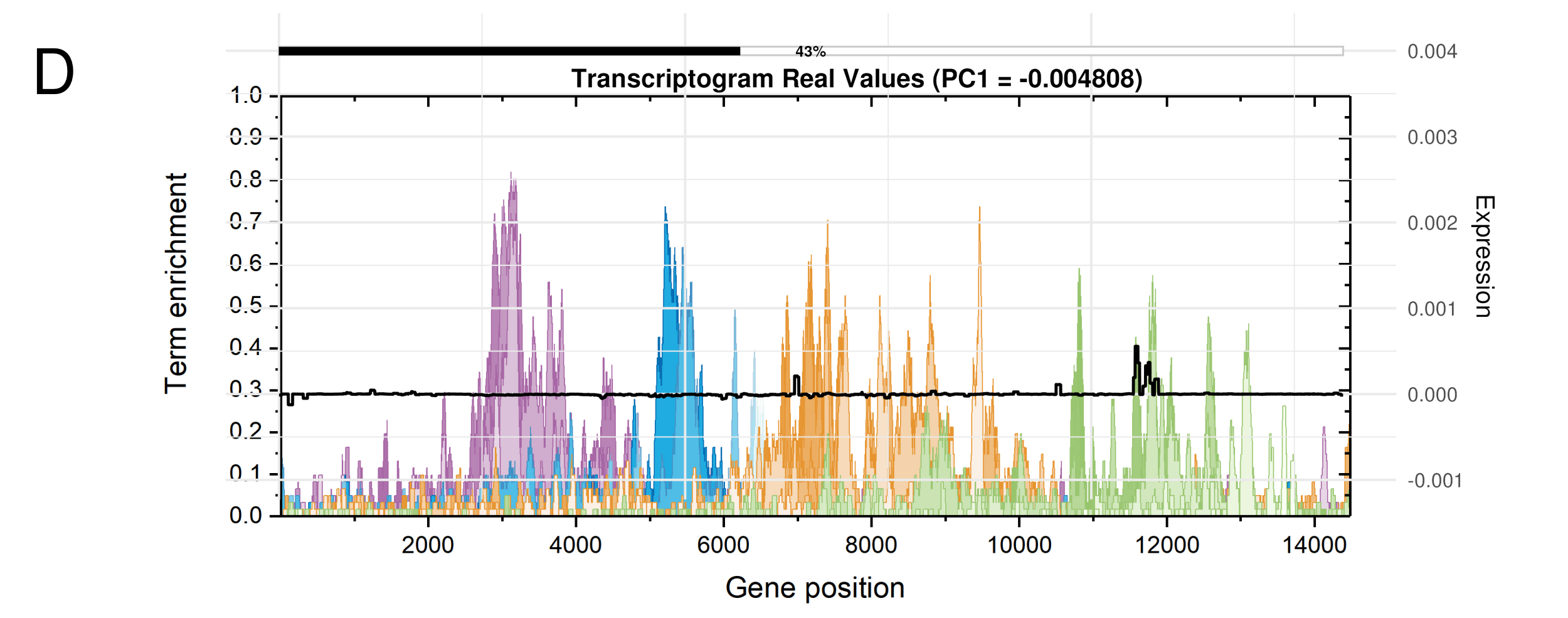

### Fig_Part_E.png

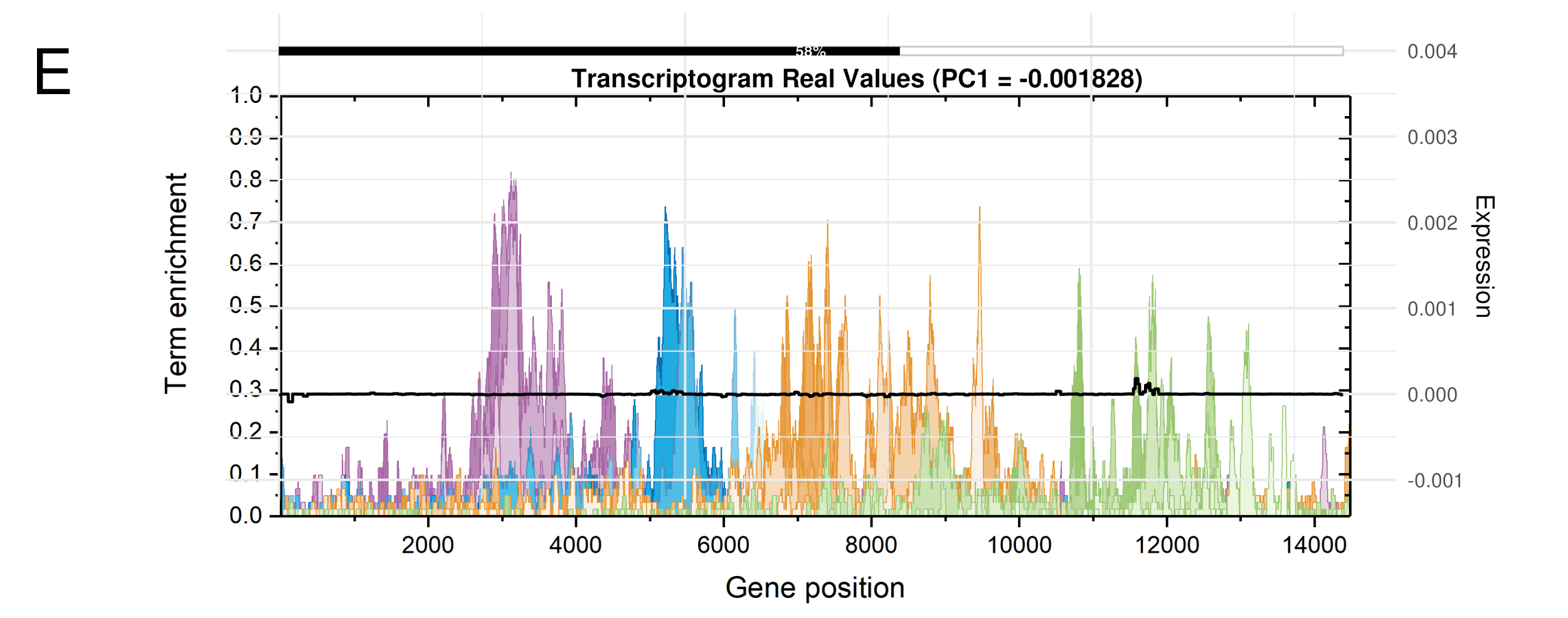

### Fig_Part_F.png

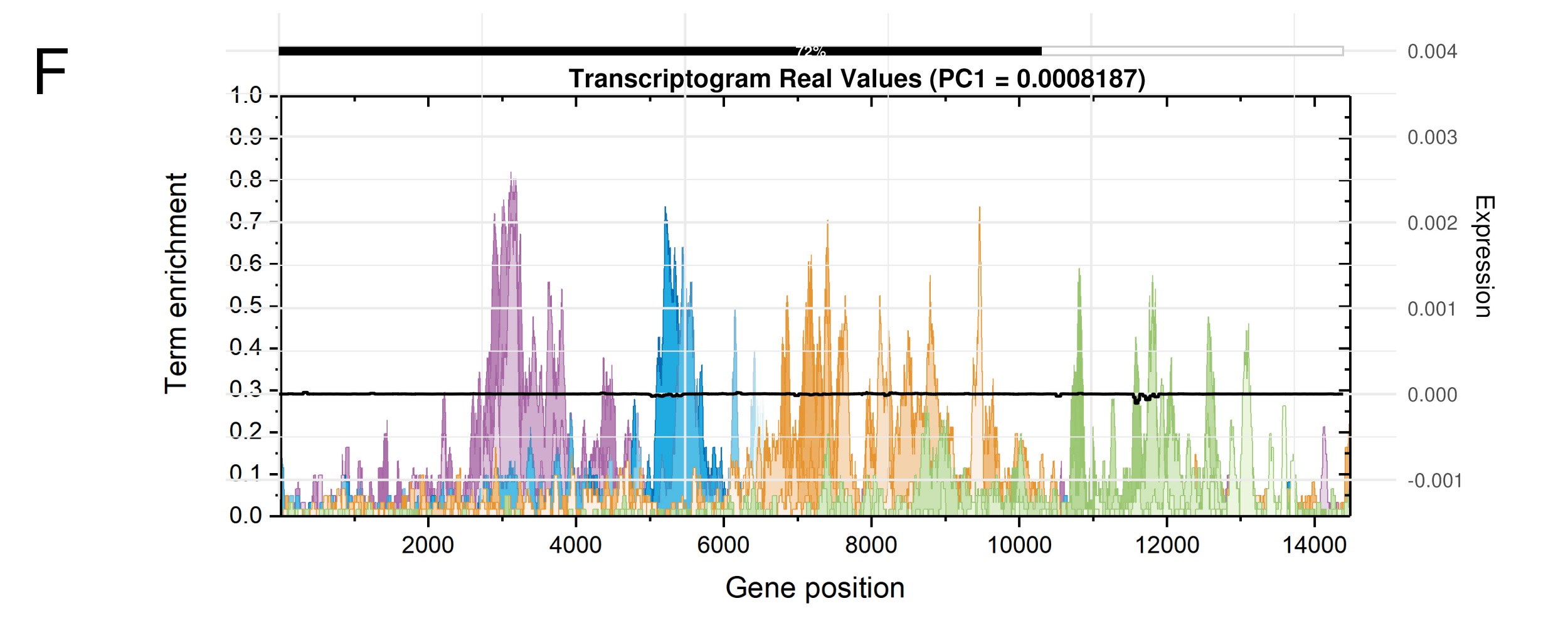

### Fig_Part_G.png

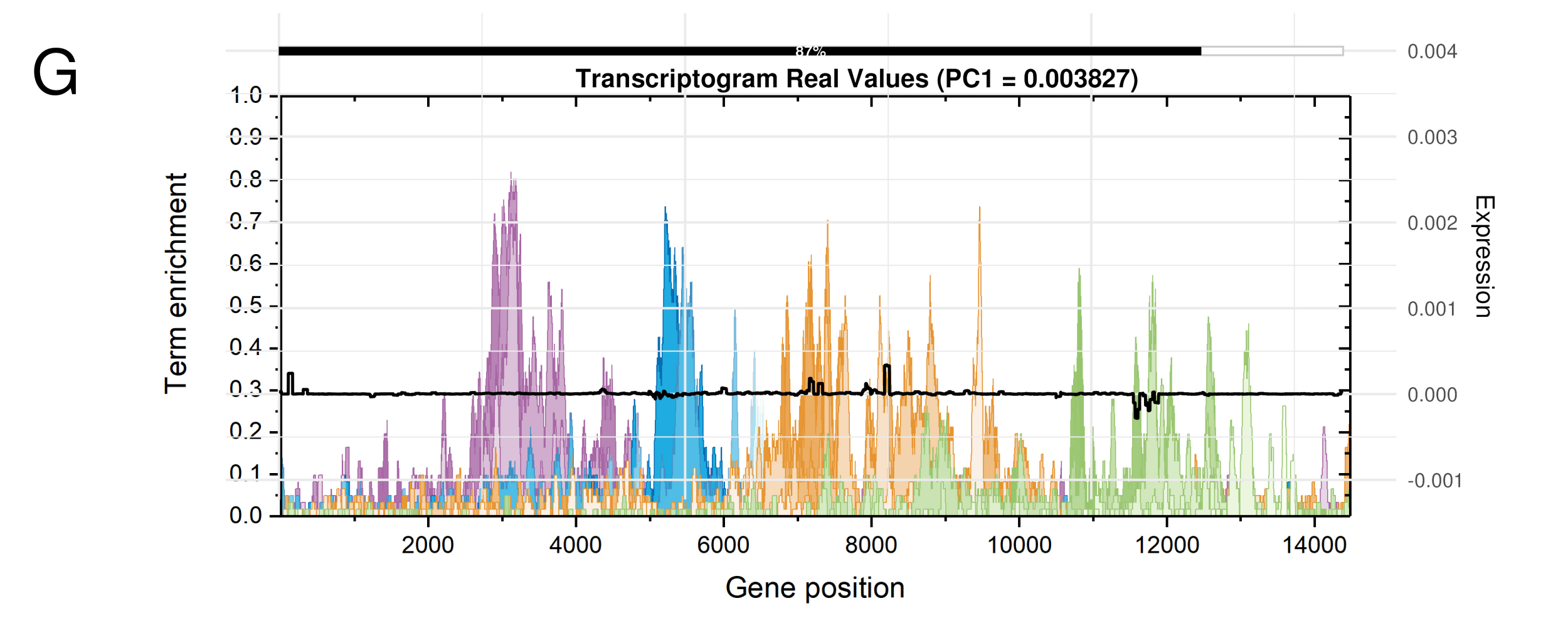

### Fig_Part_H.png

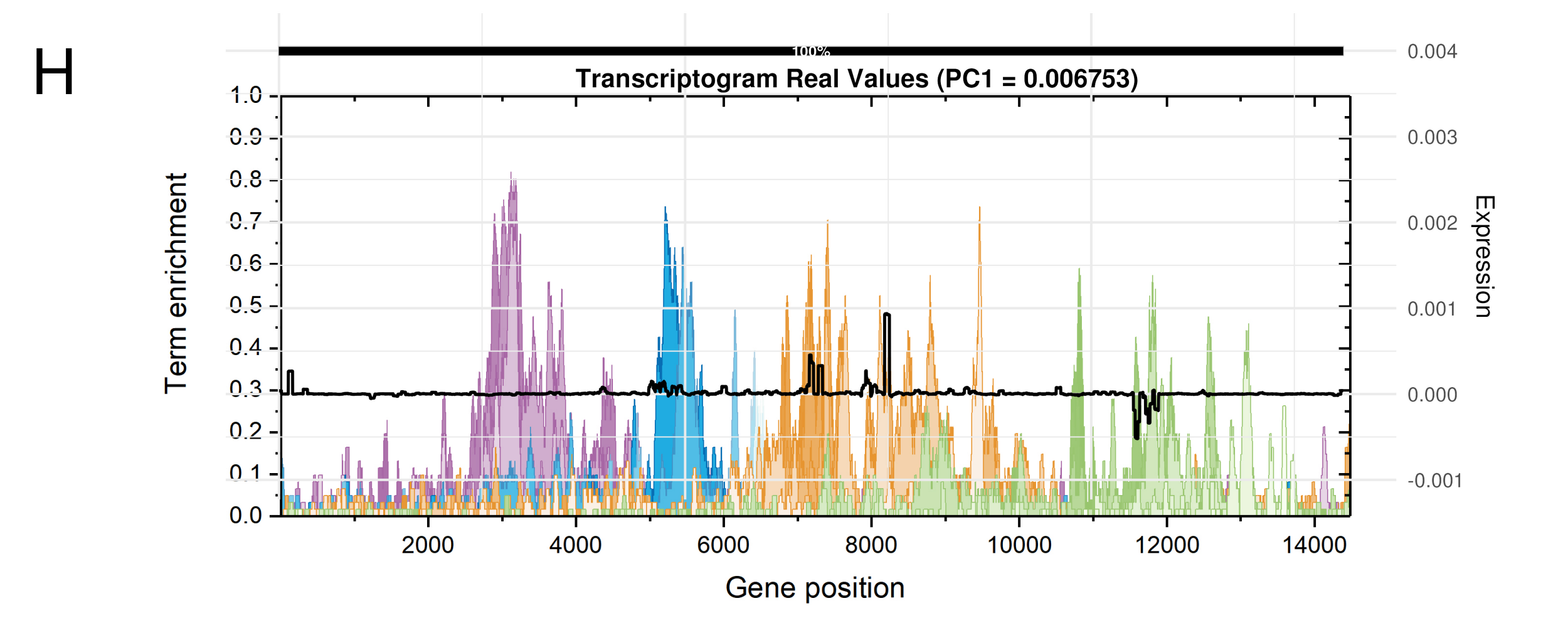

### importancia_normalizada_por_frame.png

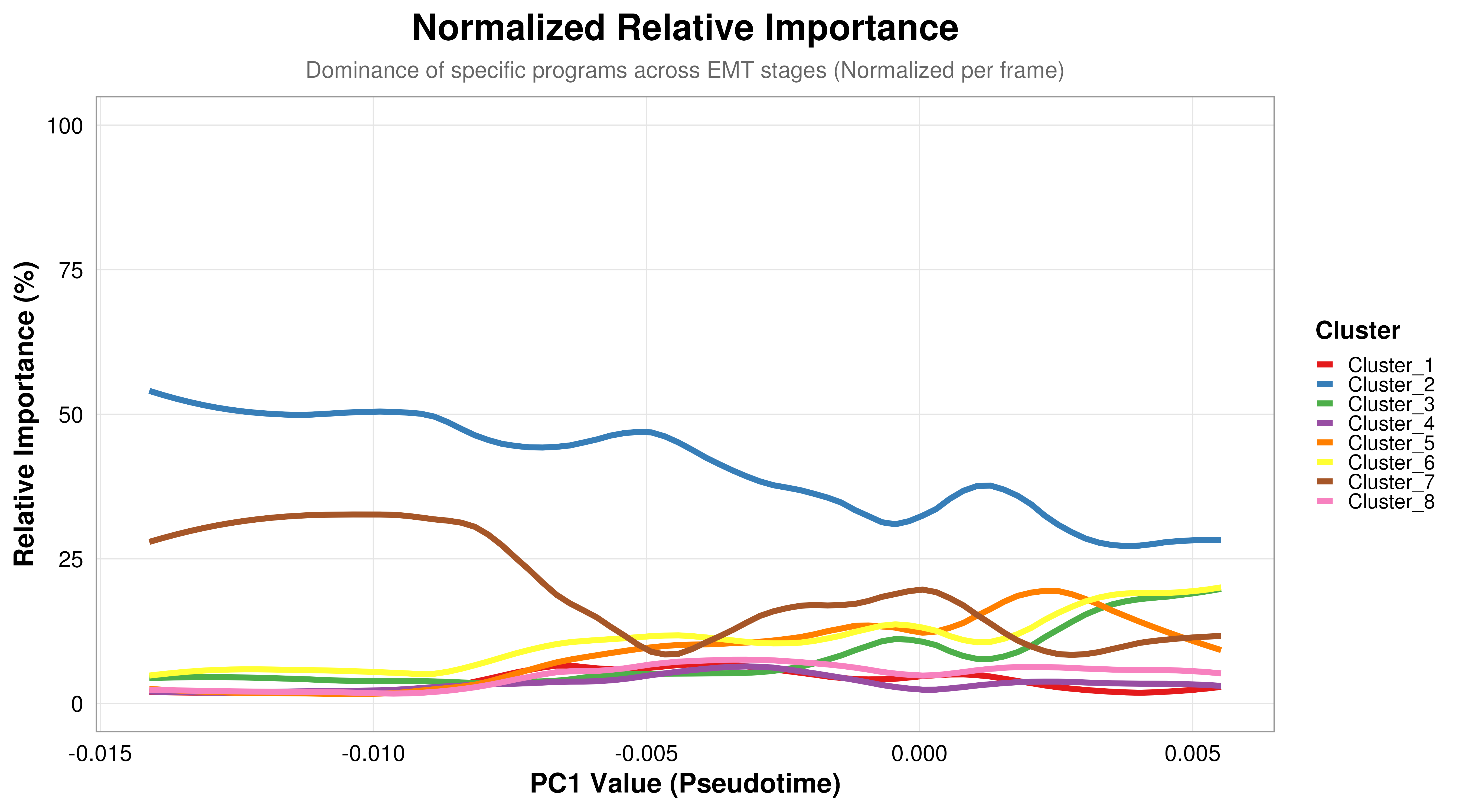

### odilon_foto.jpg

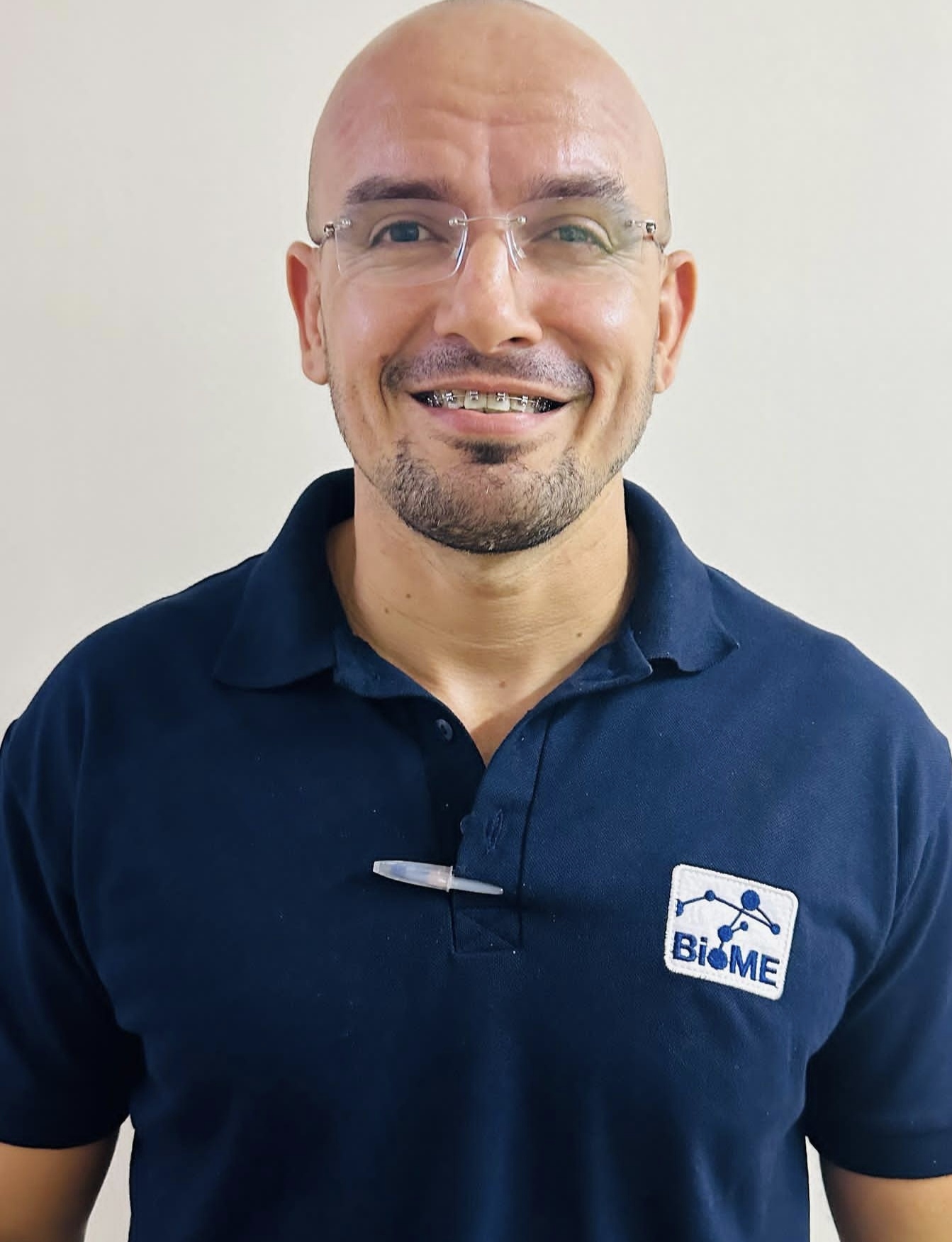

### PANEL_FINAL_ARTIGO.png

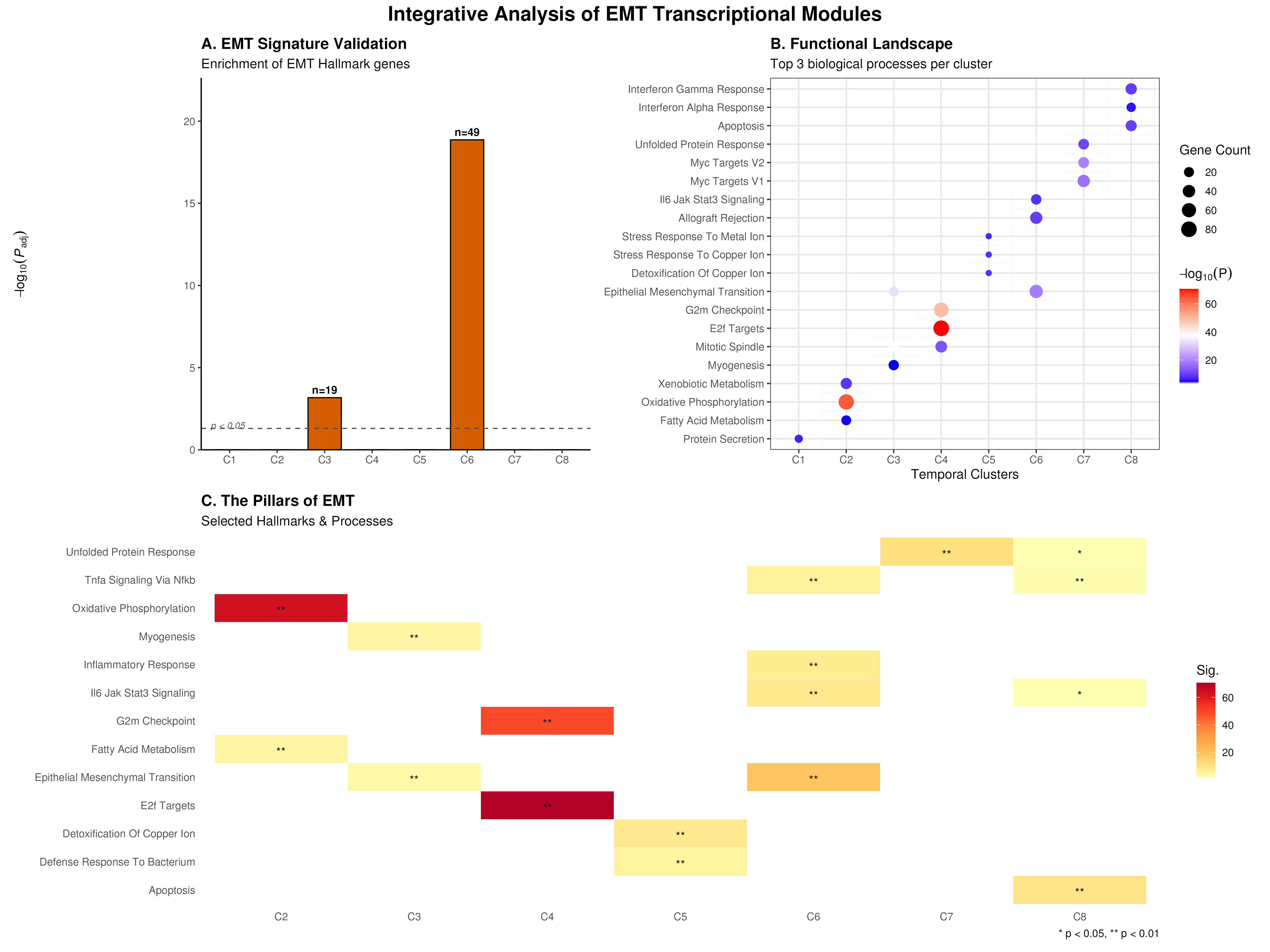

### pca_grid_PC1_vs_PC2to6_R30.pdf

#### R30 Transcriptome Trajectory (Fixed Scale)

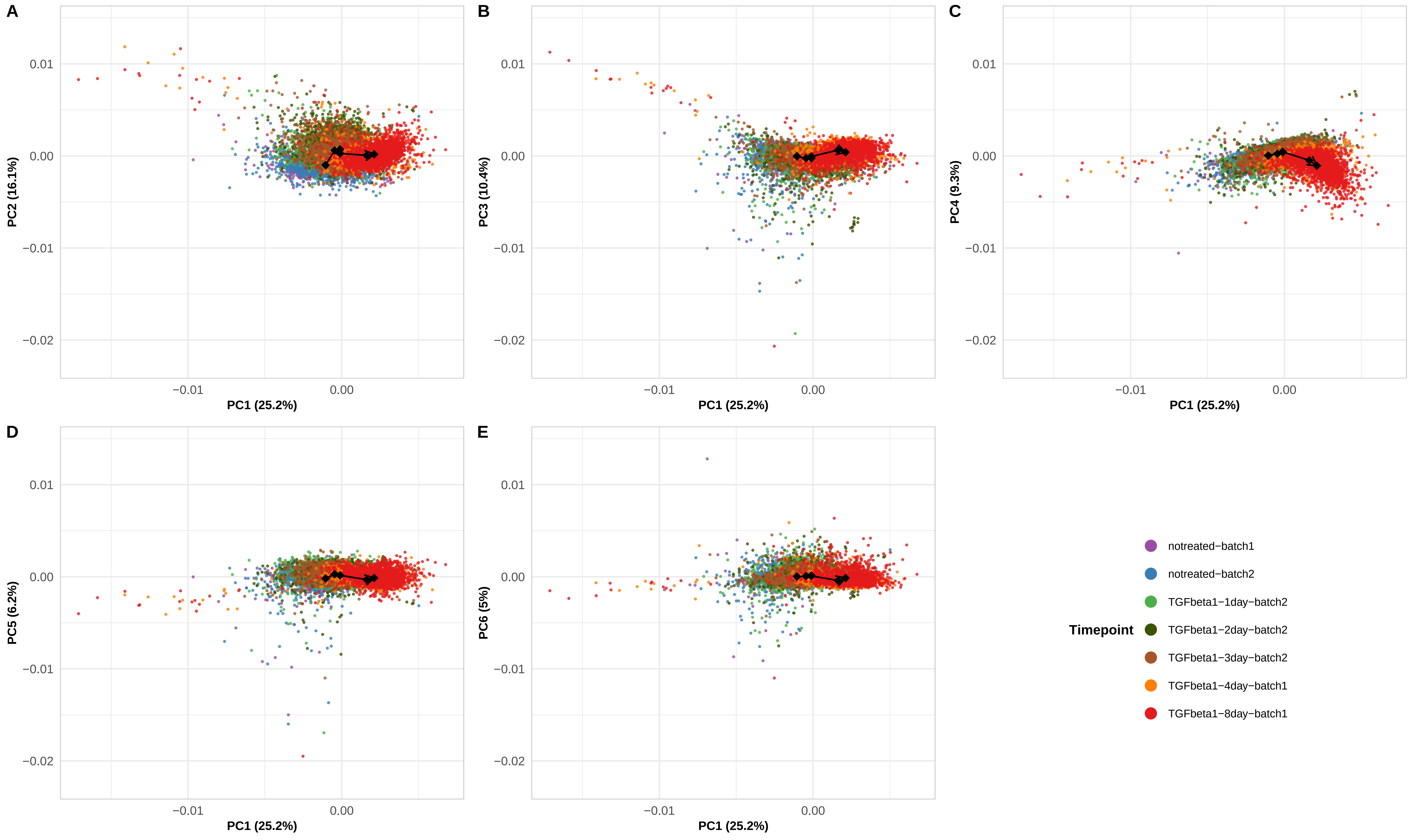

### pca_variance_complete_analysis.pdf

#### Comparative PCA Variance Analysis

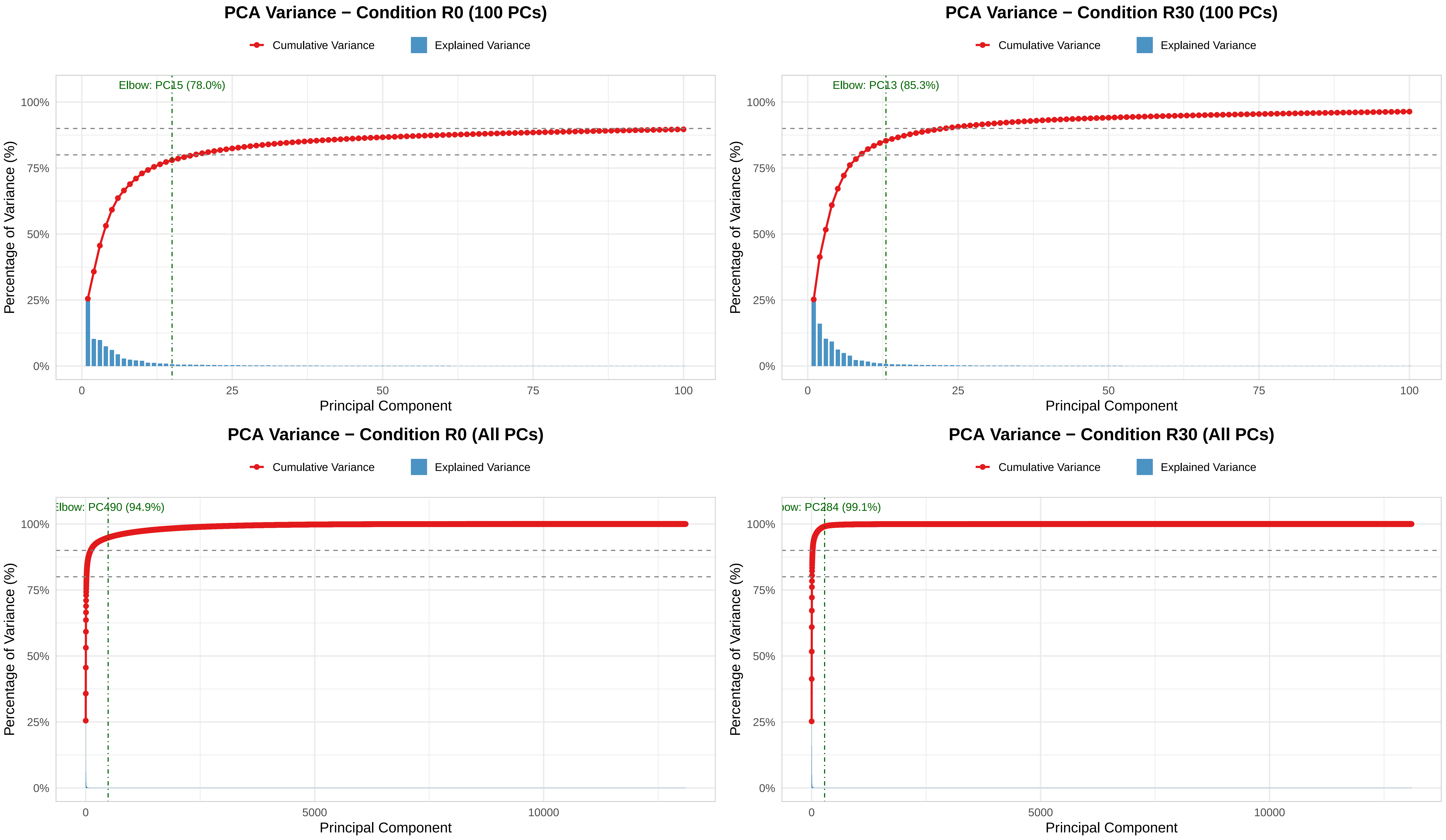

### pca_variance_R30_focused_25PC.pdf

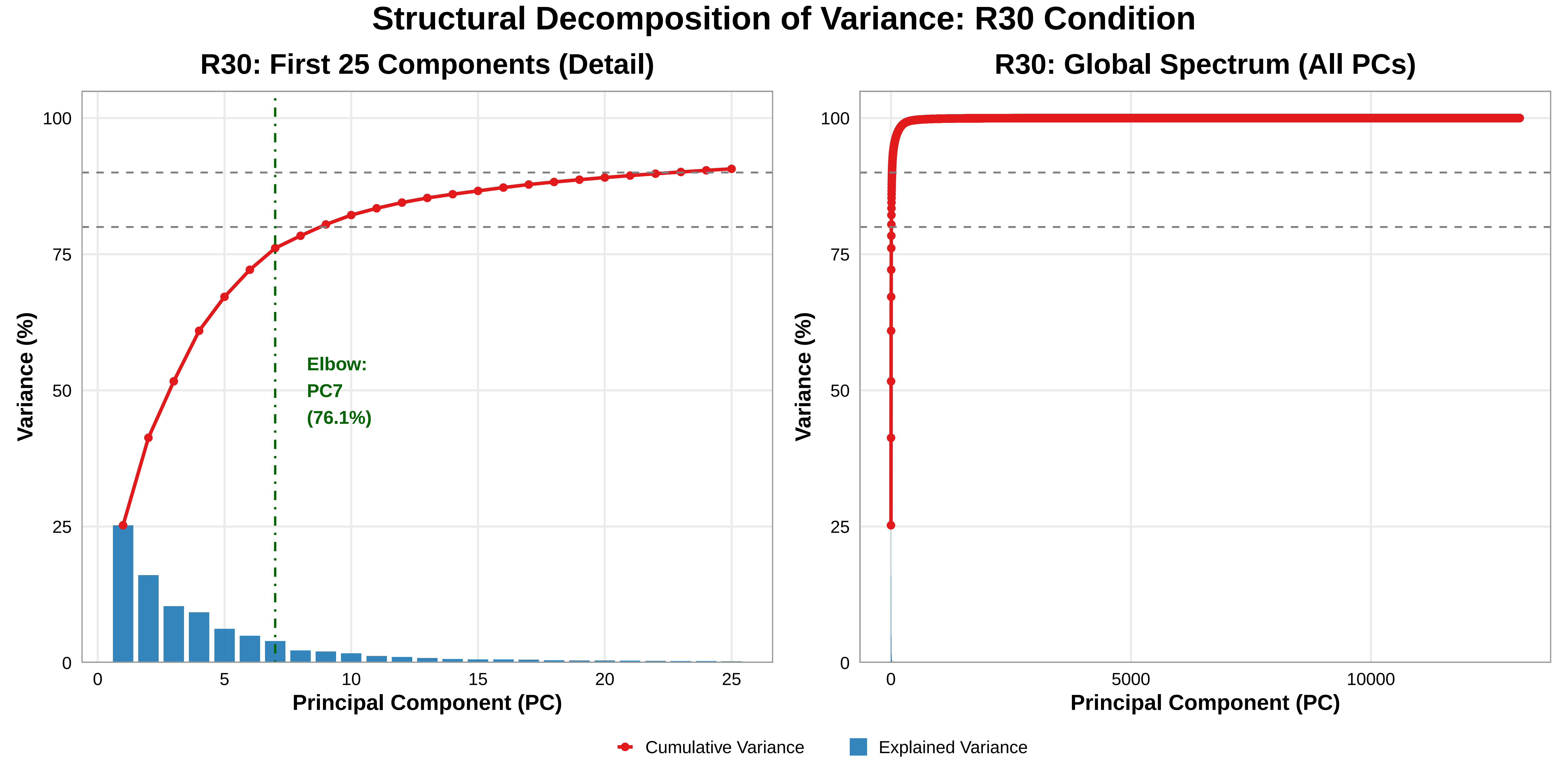

### pipeline_flowchart_new.png

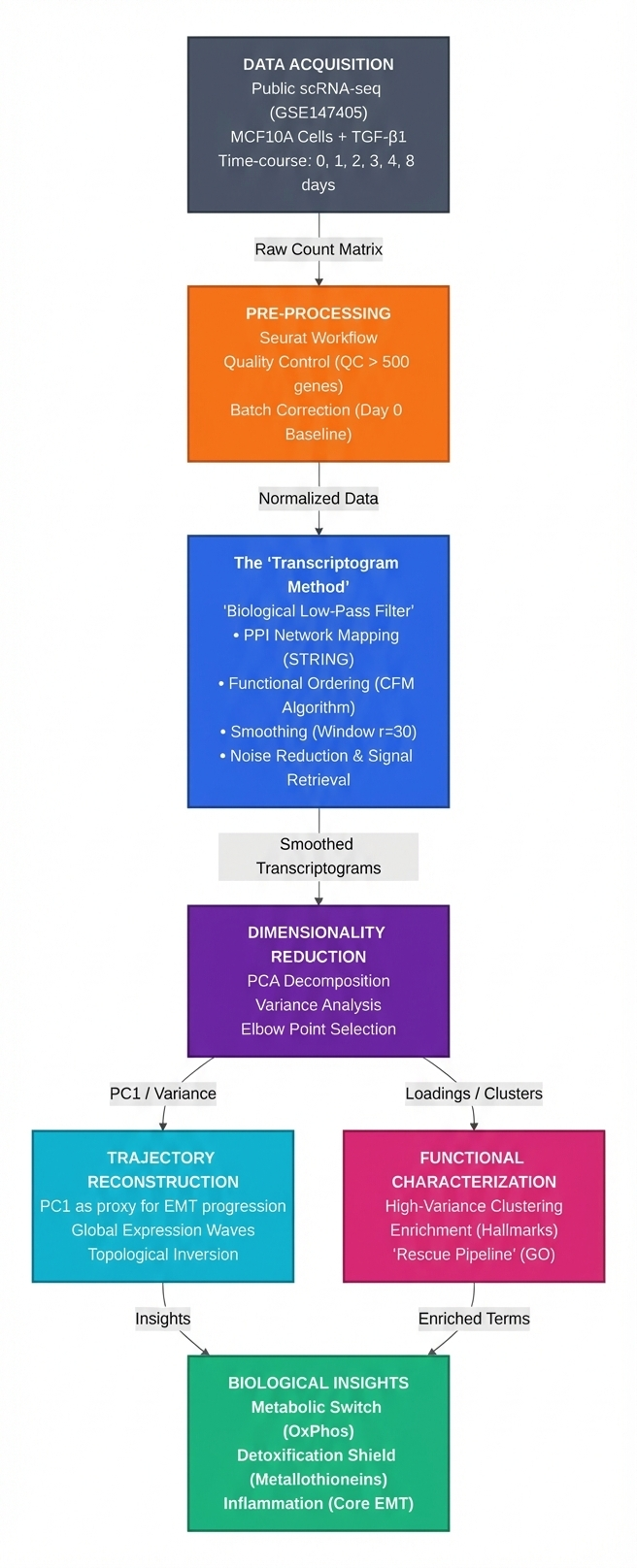

### rita_foto.png

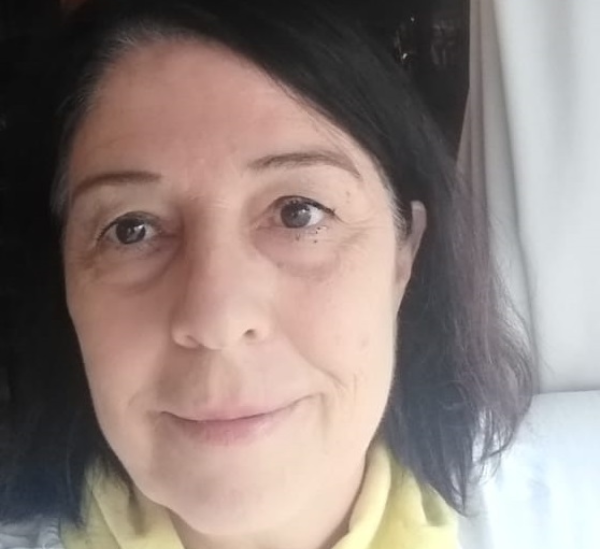

### rodrigo_foto.png

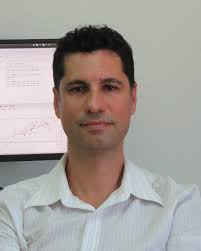

### venn_comparativo.png.png

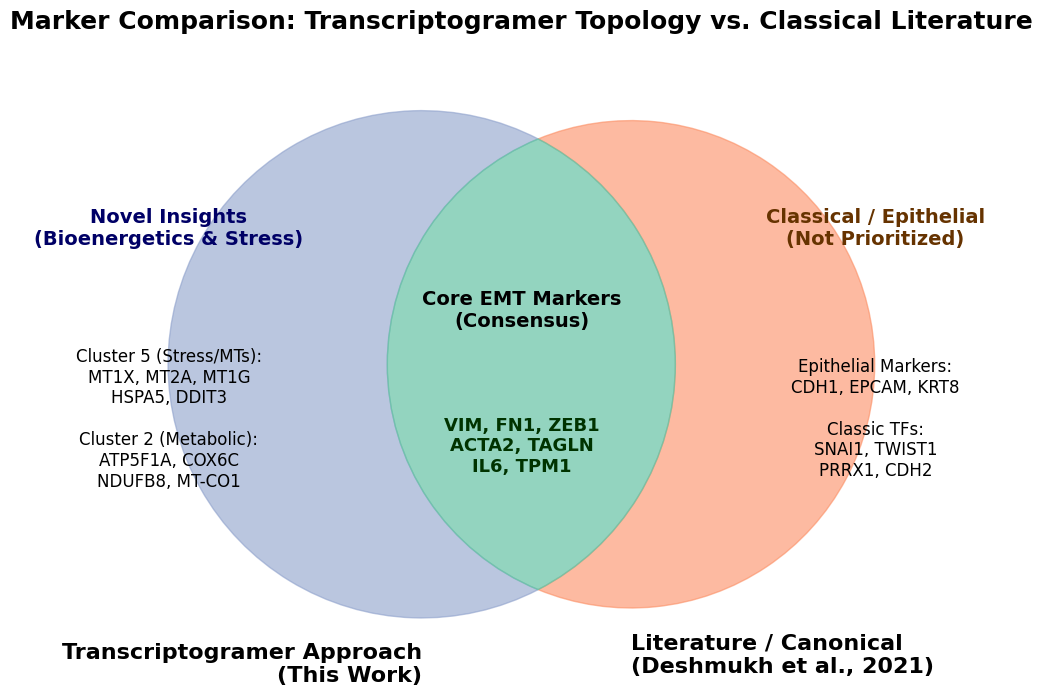
